# Exploring prophages in *Salmonella enterica*: an *in-silico* approach

**DOI:** 10.1101/2025.07.28.667176

**Authors:** F. Quelal-Madrid, C. Armijos, C. Vinueza-Burgos, L. Mejía, S. Zapata-Mena

## Abstract

Prophages play a crucial role in shaping host physiology, influencing traits such as virulence and antimicrobial resistance. In *Salmonella enterica*, prophages can constitute up to 30% of the accessory genome, contributing significantly to its genetic diversity and adaptability. In this study, we employed an *in-silico* approach to identify prophages within *S. enterica* genomes obtained poultry and clinical isolates.

A total of 151 *S. enterica* sequences were assembled using SPAdes and subsequently quality-filtered with Abyss and BUSCO. Prophage sequences were detected using Phigaro and CheckV, followed by quality assessment with PHASTEST. The identification was performed using a custom database comprising 60 *S. enterica* prophages from previous studies. Comparative genomics, including gene annotation and recombination detection, were performed on representative prophages. Virulence genes were identified using VFDB and VirulenceFinder databases. Lastly, phylogenetic analysis of the prophages was carried out based on terminase and integrase sequences.

We obtained 142 *S. enterica* genomes to analyze their prophage content. *Peduovirus pro483* was highly prevalent among *S.* Infantis isolates and carried two cargo proteins: an Imma/Irre metalloendopeptidase and a fimbrial protein. In contrast, a related *S.* Enteritidis isolate also harboring *Peduovirus pro483* carried a different set of cargo genes—HEPN-MAE-28990 and a cytosine-specific methyltransferase—suggesting horizontal transfer or a recent phage acquisition event. *Enterobacteria phage ST104*, found in *S.* Typhimurium, exhibited nearly complete identity to its reference and encoded superinfection exclusion proteins (SieA, SieB), enhancing phage resistance. Phylogenetic analyses supported their classification and revealed distinct evolutionary relationships tied to host serovar and environment.

To our knowledge, this is the first study to describe prophage diversity in *S. enterica* strains isolated from Ecuadorian poultry farms, providing insights into their potential role in strain adaptation and public health relevance.

## Introduction

Phageomes are estimated to comprise approximately 80% of the virome in some environments, particularly gut microbiomes (Dion et al., 2020; Henrot & Petit, 2022; Wendling, 2023) and can represent >20% of their host’s genome (Dion et al., 2020; Nishijima et al., 2022; Wendling, 2023). Prophages are specific forms of bacteriophages that are integrated into the bacterial genome as lysogens and remain latent until certain environmental or host physiological conditions trigger their excision (Dion et al., 2020). While residing latent in a genome, prophages can influence several aspects of their host’s physiology, such as virulence, metabolic intake, antimicrobial resistance, and phage infections, making them an interesting focus, especially for gastrointestinal tract pathogens like *Salmonella* (Nayfach et al., 2021; Tisza & Buck, 2021; Wahl et al., 2019).

In *S. enterica*, prophages constitute up to 30% of the accessory genome and are recognized as one of the main contributors to the species’ diversity (Wahl et al., 2019). Their impact has been traditionally studied through the expression of virulence genes and other characteristics that affect the adaptation of *S. enterica* to different environments, but recent studies have focused on the influence of prophages in the infection dynamics, through the expression of genes that confers resistance to certain groups of phages (Dion et al., 2020; Henrot & Petit, 2022; Wahl et al., 2019). A clear example is observed in many p22-like phages which harbor superinfection exclusion genes that prevent the infection of multiple phages by inducing abortive infection or surface antigen modification (Owen et al., 2021). Research on major serovars such as Enteritidis, Typhimurium, and Infantis has primarily focused on prophages as evolutionary markers, given their tendency to become fixed in populations. However, studies examining the specific role of prophages in shaping serovar adaptation remain scarce. It has also been observed that prophage φSE12 in *S.* Enteritidis confers specific virulence to mice, specifically through the expression of *sodCI* and *sopE* genes, which also have been observed in other prophages like *Gifsy-2* in *S.* Typhimurium conferring macrophage invasion properties and subsequent systemic infections in mice and other animal models (Araya et al, 2010). In *S.* Infantis, data regarding prophages is still scarce but there are studies linking the presence of pESI-like plasmids to specific prophage lineages like P2-like phages, helping clarify the adaptation mechanisms and the polyphyletic nature of this serovar (Gymoese et al., 2019; Andrews et al, 2024).

Understanding the influence of prophages in *S. enterica* is crucial for comprehending the emergence of certain strains or serovars. This influence operates through the expression of virulence genes, promoting the success of these strains in specific environments (Cohen et al., 2020; Gymoese et al., 2019; Moura de Sousa et al., 2021; Weissman et al., 2018). However, despite their recognized impact, the presence, distribution, and functional roles of prophages in *S. enterica* genomes remain largely unexplored. The limited number of studies addressing this aspect highlights significant knowledge gaps, particularly regarding the variability of prophage content across different serovars and its potential contribution to bacterial adaptation and pathogenicity.

This study utilized bioinformatic tools to identify *S. enterica* prophages across 142 genomes derived from poultry farms, chicken carcasses, and clinical cases. The research focused on evaluating the presence of virulence genes and examining their diversity within the *S. enterica* genomes.

## Methodology

### Genome Assembly

The genome assembly protocol utilized 151 Illumina NextSeq paired-end sequences obtained from a previous study (Mejía et al., 2020), available under bioproject PRJEB37560. Details regarding sequence names and serovars are provided in Supplementary Table 1. Trimmomatic (v0.39) (Bolger et al., 2014) was employed to trim ambiguous nucleotides from paired-end sequences, ensuring a quality threshold >20 and a minimum length of 100 bp. Quality control was performed using MultiQC (v1.16) (Ewels et al., 2016), assembly was performed using SPAdes (v3.15.4) (Prjibelski et al., 2020), employing a cut-off value of 25 and the “--careful” setting to reduce the number of mismatches and short indels. Only assembled scaffolds were utilized to ensure the identification of large prophage genomes and/or avoid fragmented prophage genomes. Assembly quality was assessed with Abyss (v2.3.5) (Jackman et al., 2017), utilizing the “abyss-fac” flag, and BUSCO (v5.5.0) (Manni et al., 2021). To guarantee less fragmented identified prophage genomes, and to obtain more comprehensive and informative results, we selected genomes meeting specific criteria: assemblies with <150 scaffolds, N50 >150 kb, L50 <15, genome completeness >85%, and a genome size >4.1 MB.

### Prophage sequences identification

Assembled genomes were analyzed for prophage sequences, which were identified using Phigaro (v2.4.0) (Starikova et al., 2020) and CheckV (v1.5) (Nayfach et al., 2021). Proviral sequences shorter than 6,000 bp were excluded, as the shortest entry in our database was 6,408 bp. All prophage sequences obtained with both tools (available at https://github.com/fran1722/Exploring-prophages-in-Salmonella-enterica-an-in-silico-approach.git) were concatenated into a single file to assess their completeness and quality using the PHASTEST online tool (Arndt et al., 2019). Briefly, prophage sequences uploaded to PHASTEST are classified based on a score: “intact” (score ≥ 90), “questionable” (score ≥ 70 and < 90), and “incomplete” (score < 70). This score is determined by the identity percentage of identified phage proteins, the number of phage features found in the query sequence, and the size of the prophage sequence (Arndt et al., 2019).

Identified prophage sequences were extracted from the *Salmonella* genomes for prophage species identification using a custom database with BLAST command line applications (v2.13.0+) (Altschul et al., 1990; Morgulis et al., 2008). The database consisted of 160 reference prophage genomes retrieved from prior studies (Gao et al., 2020; Switt et al., 2015), which established that these genomes were exclusive to or related to *S. enterica* to avoid including prophages with a wide range of hosts (Supplementary Table 2). The completeness and correct species assignment of the references were assessed through manual comparison with the NCBI Nucleotide and RefSeq databases (Sayers et al., 2022) and the latest information from the International Committee on Taxonomy of Viruses (ICTV) (Turner et al., 2021; Walker et al., 2020). Species identification was performed using blastn with default parameters. Previous studies (Ha & Denver, 2018; Hatfull et al., 2010; Pope et al., 2011) have used matches representing 45%-58% of the reference genome with > 90% of identity to evaluate gene function and completeness level, in our case, we used matches with an identity percentage and query coverage > 60% for subsequent analysis. Prophage sequences that were not identified using the custom database were further analyzed using the NCBI virus database, using the online BLAST service (Hatcher et al., 2017).

### Prophage comparative genomics

Each of the selected prophage sequences were aligned with their respective reference using MAFFT (v7.520) (Katoh et al., 2019), and subsequently, only a representative prophage sequence for each prophage species and size was selected for genome annotation.

Representative prophage sequences had their genomes annotated using Prodigal (v2.6.3) (Hyatt et al., 2010) for gene and ORF calling, specifying a bacterial-plastid genetic code. Gene sequence identification was performed using BAKTA (v1.8.1) (Schwengers et al., 2021), PHASTEST (Arndt et al., 2019), and the HmmerWeb Pfam 35.0 database (Potter et al., 2018) from EMBL. Annotated genomes were visualized and curated using Geneious Prime (v2023.0.2) (Kearse et al., 2012) to evaluate nonsense stop codons and incorrectly identified ORF*S.* Hypothetical gene sequences were aligned against the UniProt (Coudert et al., 2023) database using the BLAST online service. Curated annotated genomes (available at) were compared with their respective reference using the DigAlign (v2.0) (Nishimura et al., 2024) web service to detect recombination and moron elements in the prophage sequences. Obtained comparison graphics were curated using Adobe Illustrator for a better understanding.https://github.com/fran1722/Exploring-prophages-in-Salmonella-enterica-an-in-silico-approach.git) were compared with their respective reference using the DigAlign (v2.0) (Nishimura et al., 2024) web service to detect recombination and moron elements in the prophage sequences. Obtained comparison graphics were curated using Inkscape (v.1.4) for a better understanding.

### Prophage virulence genes identification

Additional to the genome annotation, representative prophage sequences obtained from the comparative genomics were analyzed for previously reported virulence genes using the virulence factor database (VFDB) (Liu et al., 2022) and the VirulenceFinder database from the Center of Genomic Epidemiology from the Technical University of Denmark (DTU) (Malberg Tetzschner et al., 2020). A custom database using the genes from these repositories was created using the BLAST command line applications (v2.13.0+) (Altschul et al., 1990; Morgulis et al., 2008) as previously described). https://github.com/fran1722/Exploring-prophages-in-Salmonella-enterica-an-in-silico-approach.git).

### Prophage classification and taxonomy analysis

Prophage sequences were analyzed to determine their relationships with other related prophage groups and to corroborate their taxonomic assignments. Representative prophages previously obtained from the comparative genomics sequences were grouped with other taxonomically related phages from the ICTV to assess their relationships within their corresponding prophage group (Supplementary Table 2). Subsequently, the amino acid sequences for the terminase subunits (large and small), and integrase were extracted from each prophage group, analyzed, and aligned using MAFFT (v7.520) (Katoh et al., 2019). These alignments (available at https://github.com/fran1722/Exploring-prophages-in-Salmonella-enterica-an-in-silico-approach.git) were used to construct a Neighbor-Joining tree with Geneious Prime (v2023.0.2) (Kearse et al., 2012), which was curated with FigTree (v1.4.4) (http://tree.bio.ed.ac.uk/software/figtree/). Four phylogenetic trees were constructed: one with concatenated sequences of both terminase subunits, one with integrase (Int) sequences, one with Terminase large subunit (TerL) sequences, and one with Terminase small subunit (TerS) sequences.

## Results

### *Salmonella* genomes and prophage identification

After removing duplicated or ambiguous reads from a total of 151 *S. enterica* data sets, the remaining reads were assembled into genomes. However, only 142 genomes met the required criteria for identifying unfragmented prophage genomes. These criteria included having < 150 scaffolds, an N50 > 150 kB, an L50 < 15, genome completeness > 85%, and a genome size exceeding 4.1 Mb. (Supplementary Table 1; Supplementary Figure 1).

The 142 assembled genomes exhibited an average of 152.63 scaffolds, with an N50 of 153,849 bp and an L50 of 6.62. The assembled genomes maintained an average genome size of 4.71 Mbp (Supplementary Figure 1). Among these, 136 genomes were identified as *S.* Infantis (131 from poultry farms and 4 from clinical isolates), 3 as *S.* Enteritidis (1 from a poultry farm and 2 from clinical isolates), and 3 as *S.* Typhimurium (all from poultry farms). (Supplementary Table 2).

**Figure 1.**
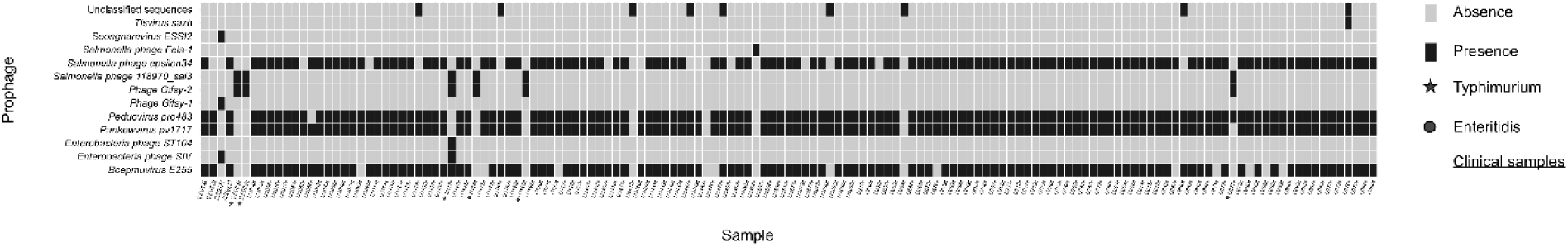
Heatmap representing the 12 distinct *Salmonella* prophage-associated sequences identified across the 142 *S. enterica* genomes.

Twelve distinct *Salmonella* prophage-associated sequences were identified with an identity ranging from 69.10% to a maximum of 99.91%, the query coverage ranged from 12% to 100% of the prophage sequence. Notably, only the *S.* Infantis genome (U1467s), did not harbor any *Salmonella* prophages. Among the 142 genomes analyzed, 45 contained five *Salmonella* prophage sequences, 71 carried four prophages, and 23 harbored three prophages. Two genomes of *S.* Infantis U1412s and U1496s had one and two prophages, respectively. (Figure 1, Supplementary table 2).

The 135 *S.* Infantis genomes showed the greatest diversity of prophage sequences detected, with a total of 9 different prophage-associated sequences. The most prevalent prophage sequences were *Bcepmuvirus E255, Pankowvirus pv1717, Peduovirus pro483*, and *Salmonella phage epsilon34*. Each of these sequences presented different sizes within the genomes ranging from 45,903 bp to only 4,198 bp. Additional prophage sequences were identified; however, these were present in only a single genome, like *Phage Gifsy-1, Salmonella phage Fels-1*, *Seongnamvirus ESSI2*, and *Tlsvirus sazh* (Table 1).

**Table 1.** Distribution of prophage sequences detected in each *S.* Infantis genome.

| ID<br>(NCBI reference<br>code) | Prophage<br>reference<br>size<br>(bp) | Phigaro +<br>checkV | *Salmonella*<br>genomes |  | BLAST |  |  | PHASTEST |  |
| --- | --- | --- | --- | --- | --- | --- | --- | --- | --- |
|  |  | Prophage<br>detected<br>size<br>(bp) | Farms<br>*n* = 131 | Clini-<br>cal<br>*n* = 4 | Percentage<br>(%)<br>ID | Query<br>Coverage<br>(%) | Max<br>Score | Total<br>Score | Completeness<br>classification |
| *Bcepμvirus E255*<br>(NC_009237) | 37446 | 17388 | 15 | 1 | 69.1 | 21 | 732 | 1467 | Intact |
|  |  | 20441 | 98 | 2 | 69.1 | 18 | 732 | 1467 | Intact |
| *Enterobacteria phage*<br>*SfV*<br>(NC_003444) | 37074 | 5826 | 0 | 1 | 76.29 | 12 | 2143 | 744 | Intact |
| *Pankowvirus pv1717*<br>(NC_011357) | 62147 | 23765 | 120 | 3 | 71.03 | 43 | 2269 | 4443 | Intact |
|  |  | 41942 | 1 | 0 | 85.94 | 36 | 2269 | 10128 | Intact |
|  |  | 42427 | 6 | 0 | 85.94 | 35 | 2269 | 9556 | Intact |
|  |  | 43182 | 1 | 0 | 85.94 | 34 | 2269 | 9556 | Intact |
| *Peduovirus pro483*<br>(NC_028943) | 29237 | 6603 | 49 | 1 | 97.47 | 99 | 11152 | 11284 | Incomplete |
|  |  | 20172* | 47 | 1 | 97.82 | 80 | 21830 | 41710 | Intact |
|  |  | 20234* | 4 | 0 | 97.82 | 81 | 21825 | 27885 | Intact |
|  |  | 31653* | 76 | 2 | 97.82 | 79 | 21830 | 41710 | Intact |
| *Phage Gifsy-1*<br>(NC_010392) | 48491 | 19209* | 0 | 1 | 99.97 | 100 | 34611 | 34937 | Intact |
| *Salmonella phage ep-*<br>*silon34* (NC_011976) | 43016 | 17887* | 115 | 3 | 94.18 | 73 | 18444 | 19774 | Intact |
| *Salmonella phage*<br>*Fels-1*<br>(NC_027984) | 42723 | 39728 | 1 | 0 | 90.16 | 46 | 16992 | 23754 | Intact |
| *Seongnamvirus ESS12*<br>(NC_047854) | 28765 | 30206* | 0 | 1 | 75.03 | 69 | 8342 | 15129 | Intact |
| *Tlsvirus sazh*<br>(NC_048065) | 49665 | 43966* | 1 | 0 | 90.36 | 87 | 27663 | 53120 | Intact |

(\*) Selected prophage lengths with an adequate identification percentage and query coverage for genome annotation and comparative genomics.

The remaining six *S. enterica* genomes included three *S.* Typhimurium, in which *Phage Gifsy-2* and *Salmonella phage 118970_sal3* were the most prevalent prophage sequences. Additionally, *Enterobacteria phage ST104* was identified as the only prophage with an integrity of 99.97% (Table 2). The other three genomes corresponded to *S.* Enteritidis, which also carried *Phage Gifsy-2* and *Salmonella phage 118970_sal3*, while one *S.* Enteritidis genome uniquely featured *Peduovirus pro483* (Table 3).

**Table 2.**
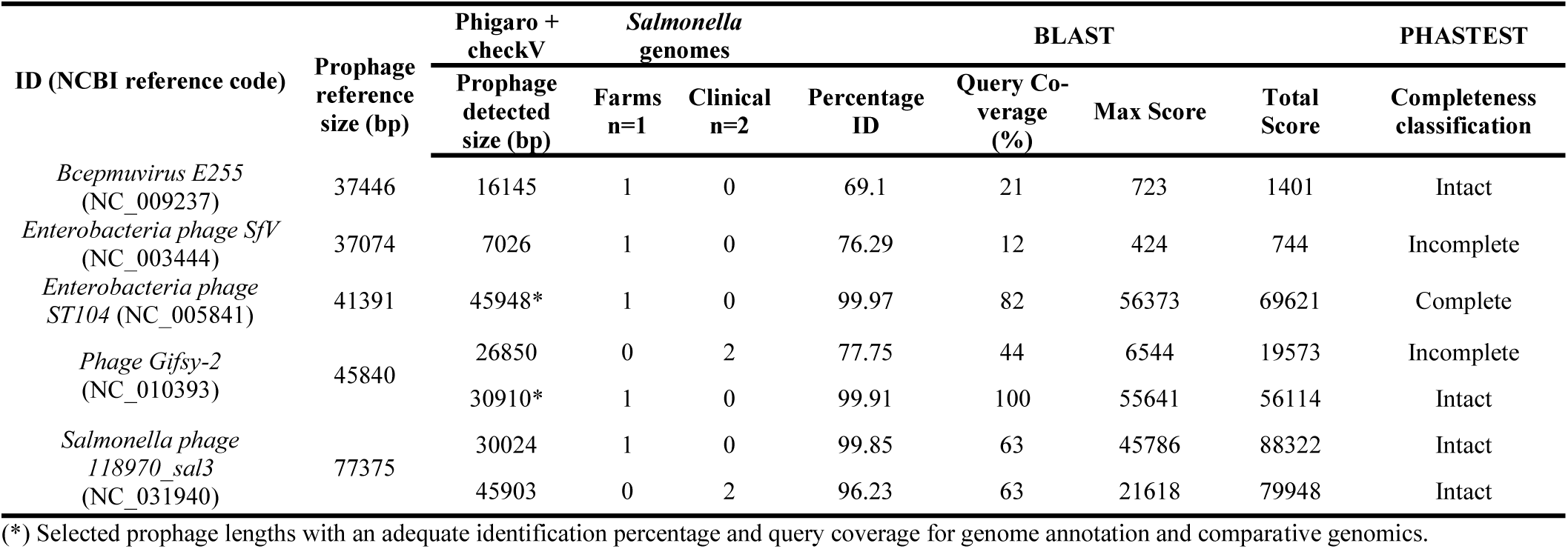
Prophage Sequences detected in *S.* Typhimurium genomes.

**Table 3.**
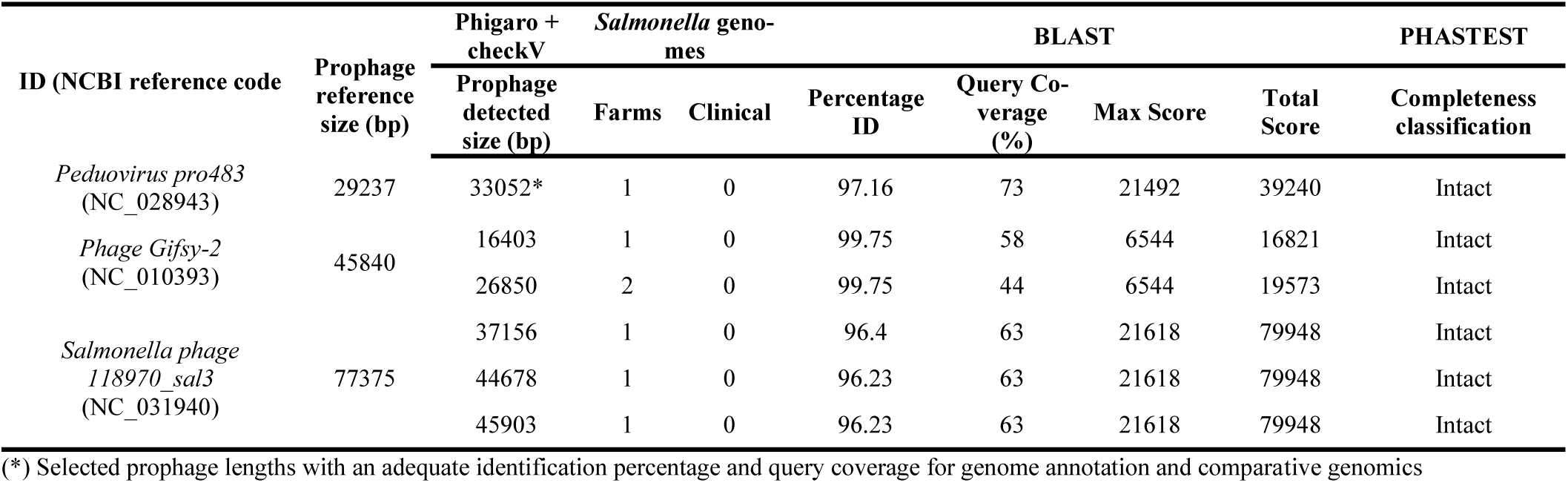
Prophage Sequences detected in *S.* Enteritidis genomes.

### *Salmonella* prophages comparative genomics and gene identification

As shown in Tables 1, 2, and 3, prophage sequences fulfilling the criteria of >95% identity and >65% query coverage were included in the analysis. A total of seven prophage sequences were compared with their respective reference genomes to identify cargo and/or virulence genes.

The different sizes associated with *Peduovirus pro483* were analyzed to identify cargo or virulence features in the representative genomes (Figure 2). We observed two main genetic blocks in common in all our *Peduovirus pro483*-like sequences with a 100% identity to the reference *Peduovirus pro483*. In all *Peduovirus pro483-like* sequences found in *S.* Infantis, regardless of size, two cargo proteins were consistently identified: Imma/Irre metalloendopeptidase and a fimbrial protein (Table 4). Interestingly, the only *Peduovirus pro483*-like sequence identified in *S.* Enteritidis shared conserved genetic blocks with those from *S.* Infantis (Figure 2). However, it carried two distinct cargo proteins: a HEPN-MAE-28990 domain-containing protein and a cytosine-specific methyltransferase (Table 4).

**Figure 2.**
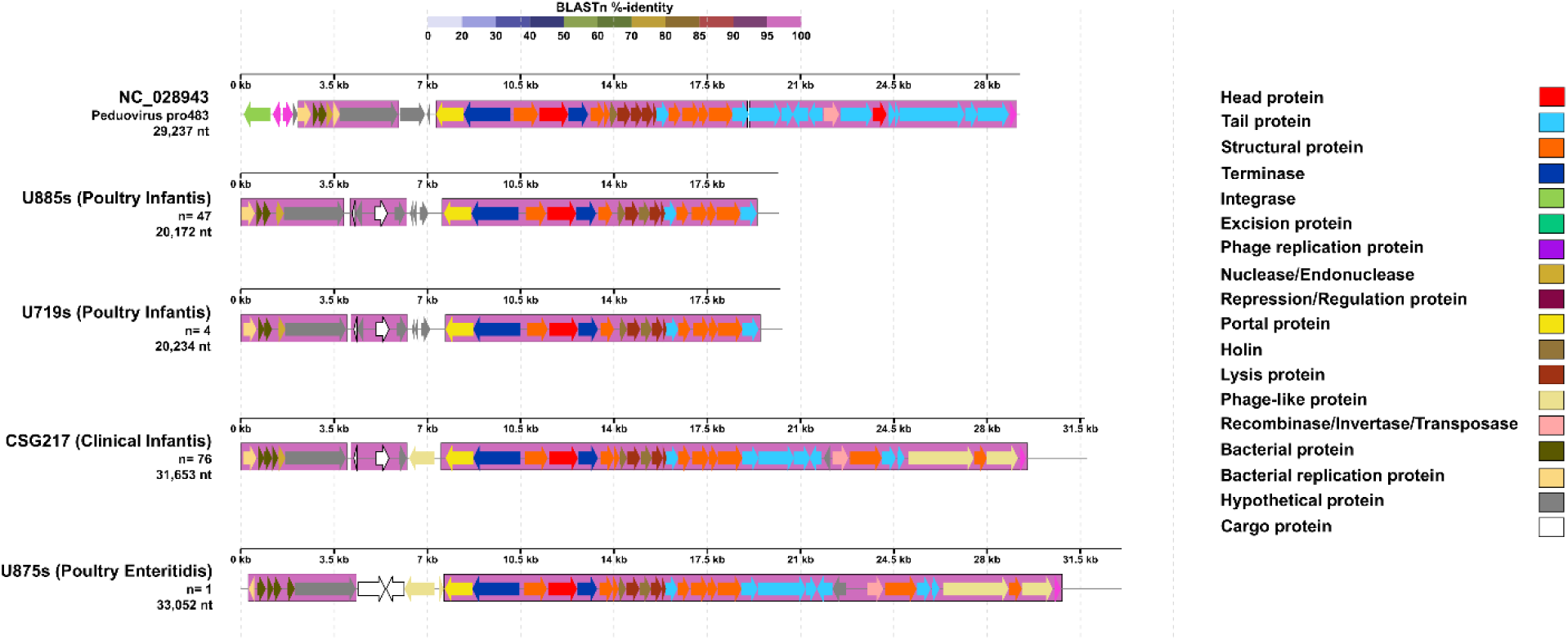
Comparative genomics graph of *Peduovirus pro483*-like sequences of varying sizes detected in *S. enterica*. The legend on the right indicates the main features identified for these prophage sequences compared to the *Peduovirus pro483* reference genome (NC_028943). At the top, a BLAST %-identity bar indicates the identity percentage of every genetic block or region compared to the reference genome, according to the color palette on the top. Obtained using DigAlign (v2.0).

**Table 4.** Cargo proteins and superinfection exclusion proteins detected in *Peduovirus pro483-like* prophage sequences.

| Protein | Genomes |
| --- | --- |
| Imma/Irre Metallo Endopeptidase | 132 (Infantis) |
| Fimbrial protein |  |
| HEPN-MAE-28990domain-containing protein | 1 (Enteritidis) |
| Cytosine-specific methyltransferase |  |

The *Enterobacteria phage ST104-*like sequence from a *S.* Typhimurium showed a high level of identity and a significant number of conserved genetic blocks compared to its reference genome (Figure 3). Almost all the *Enterobacteria phage ST104* reference genome corresponded with a 100% identity, except for three genes corresponding to virulence proteins that were not present in our sequence. While features like excisionase, and some virulence factors were absent in our detected sequence, other features like SieA and SieB, both coding for superinfection exclusion proteins, and an integrase were present (Table 5). In the same sample, a *Phage Gifsy-2*-like sequence was identified, also with a high proportion of conserved genetic blocks (Figure 4) compared to its reference genome, and carrying 2 virulence factors: *SodC*, a superoxide dismutase, and GtgA, a type III secretion system protein (Table 5).

**Figure 3.**
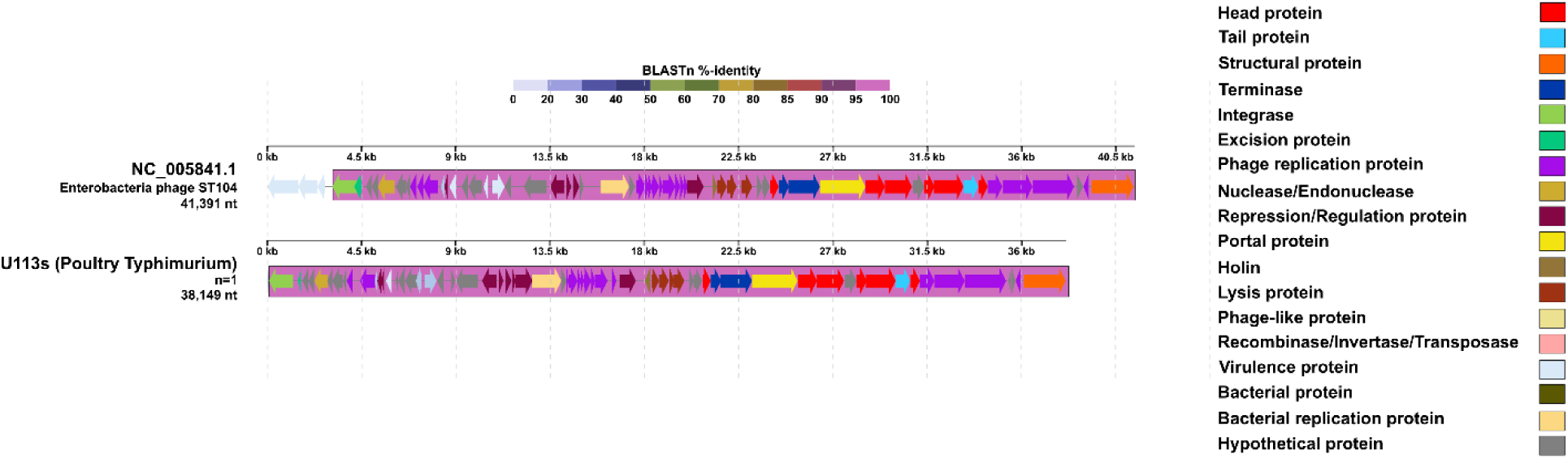
Comparative genomics graph of *Enterobacteria phage ST104-like* sequences of varying sizes detected in *S. enterica*. The legend on the right indicates the main features identified for these prophage sequences compared to the Enterobacteria phage ST104 reference genome (NC_005841). At the top, a BLAST %-identity bar indicates the identity percentage of every genetic block or region compared to the reference genome, according to the color palette on the top. Obtained using DigAlign (v2.0).

**Figure 4.**
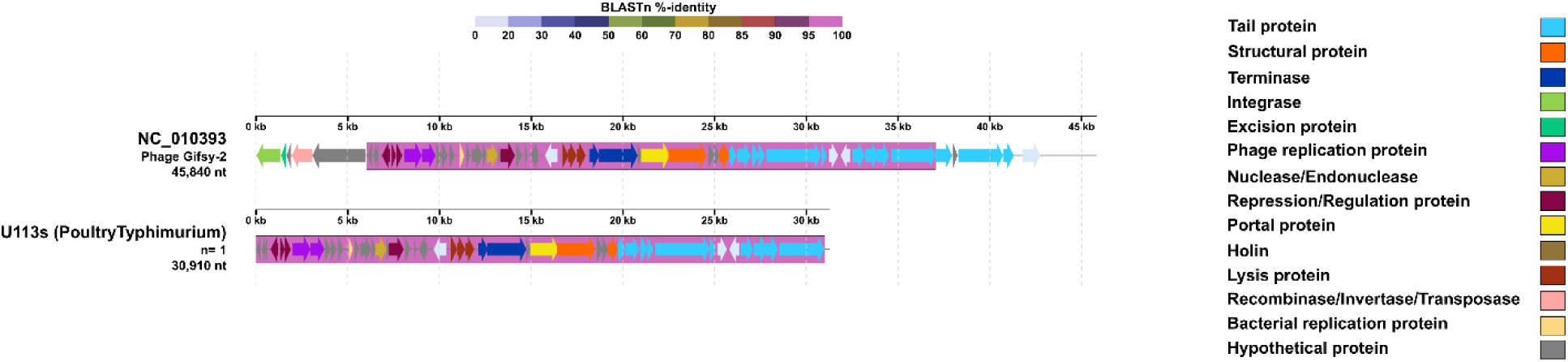
Comparative genomics of Phage Gifsy-2-like sequences of varying sizes detected in *S. enterica*. The legend on the right indicates the main features identified for these prophage sequences compared to the Phage Gifsy-2 reference genome (NC_010393). At the top, a BLAST %-identity bar indicates the identity percentage of every genetic block or region compared to the reference genome, according to the color palette on the top. Obtained using DigAlign (v2.0).

**Table 5.** Virulence proteins and superinfection exclusion proteins detected in prophage sequences in *S.* Typhimurium genome.

| Protein | Prophage |
| --- | --- |
| SieA | *Enterobacteria phage ST104-like* |
| SieB |  |
| SodC | *Phage Gifsy 2-like* |
| GtgA |  |

Prophage-like sequences for *Salmonella phage epsilon34, Phage Gifsy-1, Seongnamvirus ESSI2*, and *Tlsvirus sazh* were also compared with their respective reference genomes, but no cargo, virulence factors, or superinfection exclusion proteins were detected. Relevant features as transposase were found in *Phage Gifsy-1* and *Tlsvirus sazh,* and an integrase in *Seongnamvirus ESSI2* (Supplementary Figure 2).

To determine the taxonomic relationships of the detected prophages, we performed a phylogenetic analysis using the concatenated amino acid sequences of both terminase subunits (TerS and TerL) and the integrase (Int) of prophage sequences positive for these features (Figure 5). The resulting phylogenetic trees provided key insights into the evolutionary placement of these prophages.

**Figure 5.**
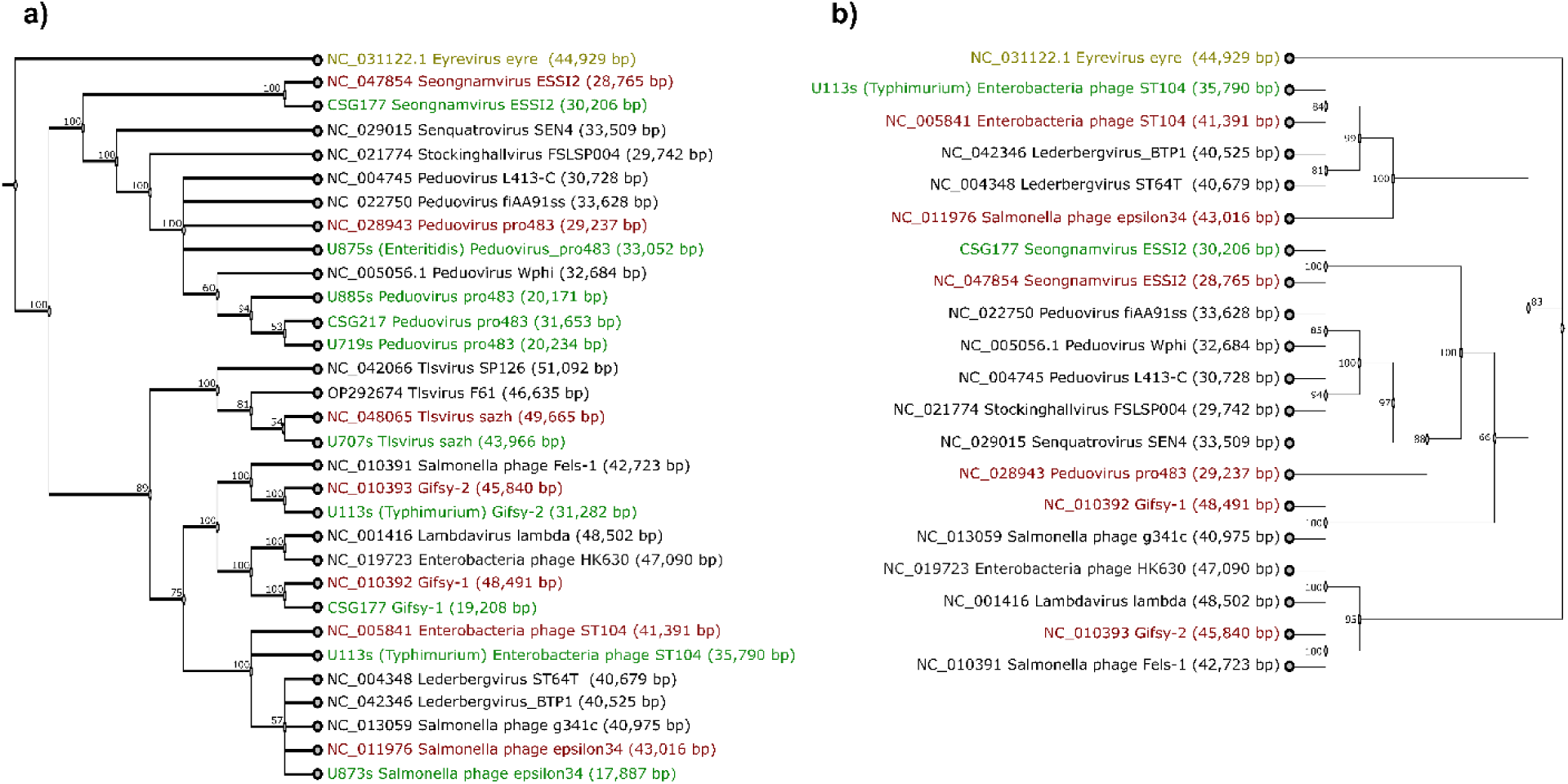
Neighbor-joining trees constructed with a) terminase small and large subunits aminoacid sequences and b) integrase aminoacid sequences. The outgroup is indicated in yellow, the prophage reference genomes are indicated in red, and the obtained prophage sequences are depicted in green. The concatenated sequences were extracted after annotation from every representative prophage sequence detected, their corresponding reference and other related prophage genomes. Obtained using FigTree (v.1.4.4)

In general terms, the phylogenetic analysis based on the terminase subunits (Figure 5a) revealed two major cluster*s.* The first cluster encompassed our *Peduovirus pro483*-like sequences along with other *Peduovirus*-related representatives, as well as our *Seongnamvirus ESSI2*-like sequence and its corresponding reference. The second major cluster was composed of three subgroups: one featuring *Lederbergvirus*-related sequences, another containing lambdoid phages that included our *Phage Gifsy-1* and *Phage Gifsy-2*-like sequences, and a third with *Tlsvirus*-related phages, including our *Tlsvirus sazh*-like sequence. Trees constructed with only the TerS (Supplementary Figure 3) and TerL (Supplementary Figure 4) sequences individually featured similar relationships as those previously described, but they showed lower support values

Similarly, the analysis based on the integrase also yielded two major clusters (Figure 5b). However, in this tree, *Phage Gifsy-2*-related sequences formed a separate group from the rest of the representatives, although the internal relationships within the other clusters remained consistent with those observed in the terminase tree (Figure 5a).

Three *Peduovirus pro483*-like sequences were found to be closely related to *Peduovirus Wphi*; all of these originated from *S.* Infantis (two from poultry farms: U885s, U719s, and one clinical isolate: CSG217). However, a *Peduovirus pro483*-like sequence detected in an *S.* Enteritidis isolate (U875s) was not related to these three specific sequences but still grouped with the rest of *Peduovirus* representatives (Figure 5a). Although we did not detect integrase in our *Peduovirus pro483*-like sequences, a similar phylogenetic relationship was observed when using integrase as a marker, but only with the reference genomes: *Senquatrovirus SEN4* and *Stockinghallvirus FSLSP004* appeared more closely related to the other *Peduovirus* phages than *Peduovirus pro483*, yet they still clustered together (Figure 5b).

Our *Enterobacteria phage ST104*-like sequence was found to be identical to its reference. It clustered with other *Lederbergvirus*-related phages, including our *Salmonella phage epsilon34*-like sequence, though this grouping had a low support value (Figure 5a). A similar relationship was observed in the integrase tree (Figure 5b); however, in this instance, *Salmonella phage epsilon34* appeared less related to the *Lederbergvirus* group, despite still clustering with them.

## Discussion

*Salmonella enterica* poses a significant public health concern due to its ability to adapt to diverse environments. This adaptability has been linked to the expression of virulence factors and other genetic traits. Recent studies have highlighted the role of prophages in modulating infection dynamics by encoding genes that confer resistance to certain phage groups (Dion et al., 2020; Henrot & Petit, 2022). Here, we investigated the presence of prophages in *S. enterica* isolates from poultry farms, chicken carcasses, and clinical sources.

Prophages are well-known for influencing host survival by providing virulence factors that enhance colonization or genes that confer resistance to environmental stress (Dion et al., 2020; Henrot & Petit, 2022; Wahl et al., 2019). While these features offer clear advantages to bacterial fitness, they can be easily overcome or lost. This loss is often due to bacterial defence mechanisms (Moura de Sousa et al., 2021; Weissman et al., 2018) and/or the influence of other mobile genetic elements (Bobay et al., 2014; Cohen et al., 2020; Gymoese et al., 2019), which can lead to defective prophages or highly fragmented prophage genomes over time (Wahl et al., 2019). This fragmentation was particularly evident in *Peduovirus pro483* and *Gifsy-2* (Figure 2 and Figure 4). Despite their high identity values, these prophages showed a loss of features like integrase and excisionase proteins, essential for their integration into and/or excision from the bacterial host. Additionally, the influence of other mobile genetic elements on the prophage genome was also evident in our *Tlsvirus sazh*-like sequence (Supplementary Figure 2). This Drexlerviridae phage, known to be strictly lytic (Crossland et al., 2019; Koonjan et al., 2020), was reported as a prophage in a single *S.* Infantis genome. Its annotation indicated the presence of a transposase, likely due to the influence of these other mobile genetic elements. This is important to consider since the analysed *Salmonella* genomes were presented with pESI-like plasmids that could be influencing the distribution and state of prophages within the bacterial genome (Mejia et al., 2020).

Despite the significant fragmentation previously described in the prophage genomes, our identification process revealed seven prophages with an identity and query coverage exceeding 60%. Among these, three—*Enterobacteria phage ST104, Peduovirus pro483*, and *Gifsy-2*—were particularly notable due to their high similarity to the reference genomes (approximately 99%, 97%, and 99% identity, respectively). These strong matches suggest a potentially greater functional relevance, particularly in influencing *Salmonella*–host interactions (Dion et al., 2020; Henrot & Petit, 2022). The presence of *Gifsy-2*, for example, is consistent with previous studies linking this prophage to virulence traits in *Salmonella* (Wahl et al., 2019), thereby reinforcing the idea that specific prophages may enhance the pathogenic potential of *S.* Infantis in poultry-related environments. Furthermore, this suggests that selective pressures are in action, maintaining incomplete or defective prophages causing phage mosaicism, where gene blocks are conserved despite different evolutionary histories (Jonge et al., 2019; Yu et al., 2017), as observed in the different *Peduovirus pro483*-like sequences extensively found in our *S.* Infantis genomes. This phenomenon can potentially be used as markers of specific strains related to specific environments (Bobay et al. 2014).

Among our *S.* Infantis genomes, *Peduovirus pro483* was the most prevalent prophage sequence, suggesting a probable adaptation of this prophage to this serovar and its current environment given the significant role prophages play in *S. enterica* genomes, especially in the adaptation to specific serovars and environments (D’Alessandro et al., 2018; Switt et al., 2015, Wahl, 2019). This type of adaptation has been reported in detail for various members of the *Peduovirus* family infecting *Escherichia coli,* especially those associated to clinical environments and Shiga toxin-producing *E. coli* (STEC), and other pathogenic strains like *E. coli* O157:H7 or O145 (Pacífico et al., 2019; Shridhar et al., 2019). *S.* Infantis have also been recognized as the fourth most prevalent non-typhoidal serovar associated with human infections worldwide (Montoro-Dasi et al., 2023), with a prevalence of 83% to 98% in broilers and poultry farms from Ecuador and Peru (Vallejos-Sánchez et al., 2019; Vinueza-Burgos et al., 2019). The high prevalence of *Peduovirus pro483* in *S.* Infantis suggests that this prophage may have specifically adapted to this successful serovar within the poultry farm environment. *Peduovirus pro483* was originally isolated from an avian pathogenic *E. coli* and *S.* Typhimurium (Gymoese et al., 2019; Petrovska et al., 2016) but also reported naturally occurring in clinical samples of *E. coli* associated with other P2-like phages like *Peduovirus fiAA91ss* and *Peduovirus Wphi* (Kondo et al., 2021; Pacífico et al., 2019), and carrying antimicrobial resistance genes (Rusconi et al., 2016). This suggests a clinical origin for the acquisition of *Peduovirus pro483* in *S.* Infantis, but more data from clinical settings is needed to have a strong confirmation, since only six *Salmonella* genomes were obtained from these settings, and only three of these carried *Peduovirus pro483.* The phylogenetic trees based on both amino acid sequences of TerL and TerS, confirmed that *Peduovirus pro483* was indeed related to other *p2*-like phages like *Peduovirus Wphi* and *Peduovirus fiAA91ss* which are found in *E. coli* genomes associated to clinical environments, as stated by Kondo et al. (2021) and Pacífico et al. (2019).

Cohen et al. (2020) have reported *Peduovirus pro483* associated with *S.* Infantis carrying pESI-like plasmids similarly to Mejia et al. (2020), but their analysis focused on genetic diversity and did not elaborate further on the genomic content of *Peduovirus pro483* or its origin. We delve deeper with our findings on *S.* Infantis by providing data on an *S.* Enteritidis genome also carrying *Peduovirus pro483*. While most of our samples were from poultry farms and chicken carcasses, 76 *S.* Infantis (including the previously mentioned 3 clinical samples) and a poultry *S.* Enteritidis presented a larger and more complete *Peduovirus pro483* genome. This could represent a novel adaptation of this prophage to *S.* Infantis, given the similar origin of our genomes. This adaptation could be driven by two possible scenarios: selective pressures within the poultry farm environment, favoring prophage variants that enhance *S.* Infantis fitness (Gao et al., 2020; Mottawea et al., 2018), or a recent acquisition of *Peduovirus pro483* given its integrity and maintenance of certain features (Bobay et al., 2014; Henrot & Petit, 2022; Wahl, 2019). Further studies are needed to identify the most likely scenario.

*Peduovirus pro483* has typically been reported carrying a bacterial host epithelial cell invasion protein called SopE, similar to other *Peduovirus P2*-like prophages found in emerging clones from poultry environments (Dziva et al., 2013; Gymoese et al., 2019; Petrovska et al., 2016; Rusconi et al., 2016). However, we found that none of the *Peduovirus* pro483-harboring genomes contained SopE. Instead, all *S.* Infantis strains with this prophage carried a fimbrial protein, a virulence factor that enhances surface adhesion and colonization (Huehn et al., 2010; Ksibi et al., 2022) and is linked to pESI-like megaplasmids (Kürekci et al., 2020; Lee et al., 2021). Interestingly, the only *S. Enteritidis* genome carrying *Peduovirus pro483* did not have this fimbrial protein in its gene annotation. Since *S. Infantis* has become the dominant serovar in poultry farms due to *S. Enteritidis* and *S. Typhimurium* vaccination (Montoro-Dasi et al., 2023), further data on these serovars are needed to confirm this observation.

The presence of *Enterobacteria phage ST104* and *Phage Gifsy-2* in our *S.* Typhimurium isolates aligns with previous reports of these prophages in this serovar (Ho & Slauch, 2000; Gao et al., 2020; Parker et al., 2021; Vaid et al., 2021). *Enterobacteria phage ST104* has been particularly associated with *S.* Typhimurium strain DT104 and related strains in cattle and beef production environments (Parker et al., 2021). Its close relationship with *p22*-like phages such as *Lederbergvirus BTP1* and *Lederbergvirus ST64T* (Colavecchio et al., 2017; Vaid et al., 2021) has made it prevalent in poultry farms, especially in serovars Typhimurium, Pullorum, and Gallinarum (Vaid et al., 2021). This prophage confers virulence factors and has also been linked to antimicrobial resistance genes (Colavecchio et al., 2017; Mohammed et al., 2023). Although our findings did not report any antimicrobial resistance genes, we did find virulence factors associated with the *Enterobacteria phage ST104* in our *S.* Typhimurium, which is supported by the association with other Lederbergvirus-like phages as stated by Vaid et al. (2021) and seen in our phylogenetic trees.

On the other hand, *Phage Gifsy-2* has also been reported as a prevalent prophage in *S. enterica*, particularly in serovar Typhimurium (Ho & Slauch et al., 2000). In general, *Phage Gifsy-2* has been reported as a lambda prophage with the ability to induce excision of other prophages and enhancing virulence in *Salmonella* hosts (Federici et al., 2020; Owen et al., 2020), which has enhanced its fixation in the genome of many *S.* Typhimurium and *S.* Enteritidis strains, especially within their pathogenicity islands (Foley et al., 2013; Liu et al., 2022; Chen et al., 2024). This aligns with our findings since our *S.* Typhimurium and *S.* Enteritidis isolates carried this prophage, and it was also associated with other lambda phages as observed in the phylogenetic trees.

Genes influencing *S. enterica* physiology through virulence factors and immunity to phage superinfection have also been found in *S.* Typhimurium. These genes were commonly carried by *Enterobacteria phage ST104* and *Phage Gifsy-2* when integrated in the same genome, which is congruent to our findings (Emmanuel, 2021; Parker et al., 2021). While avian-related *S.* Typhimurium is known to also harbor SopE in various emerging clones with *Enterobacteria phage ST104* (Kirkwood et al., 2021), our *S.* Typhimurium did not present this virulence factor. *Enterobacteria phage ST104* featured superinfection exclusion proteins like SieA and SieB known to protect from superinfection of *p22*-like and lamboid phages (Berngruber et al., 2010; Folimonova, 2012; Monteiro et al., 2019). *Gifsy-2* showed GtgA, a surface antigen modifier protein, and SodC, a virulence factor which protects *S. enterica* from oxidative stress from macrophages during intracellular infection (Ammendola et al., 2005; Figueroa-Bossi & Bossi, 1999; Golubeva & Slauch, 2006). This was confirmed in the phylogenetic trees where *Enterobacteria phage ST104* clustered among *Lederbergvirus*-like phages, such as *Lederbergvirus BTP1*, known to inhibit infection from non-*Lederbergvirus*-like phages through superinfection exclusion proteins like SieA and SieB (Owen et al., 2021), including other *p22*-like phages as previously explained.

Our phylogenetic analyses confirmed the associations of all prophages identified in this study. Based on the concatenated terminase subunits, we recovered two major clades: one comprising all *Peduovirus*-related phages along with *Seongnamvirus ESSI2*, and another including *Tlsvirus*-related phages, lambdoid phages, and *Lederbergvirus*-related phages. In the first clade, the clustering of *Peduovirus* and *Seongnamvirus ESSI2* was consistent with the current ICTV classification (Turner et al., 2021; Walker et al., 2020), as both belong to the family Peduoviridae, a relationship supported by both terminase and integrase phylogenies. The second clade included a *Tlsvirus*-related subgroup, which was absent in the integrase phylogeny due to the strictly lytic nature of these phages. However, their placement in the terminase tree was congruent with previous findings, particularly the close relationship with *Tlsvirus F61* and *Tlsvirus SP126*—phages previously reported to infect *S.* Infantis from poultry farms (Parra et al., 2023).

Other phylogenetic relationships based on the analysis of TerL and TerS sequences individually showed lower support for the relationships previously described compared to the concatenated approach. This discrepancy likely reflects the inherent challenges in viral phylogenetics, given the high diversity, modular genome architecture, and variations between species depending on their host and environment (Dion et al., 2020; Lima-Mendez et al., 2008; Mavrich & Hatfull, 2017; Rohwer & Edwards, 2002). The recommended approach for robust phage identification and classification involves using complete genome sequences and classifying them by clusters of proteins associated with viral features (Dion et al., 2020; Lima-Mendez et al., 2008; Maffei et al., 2021). However, we discarded this approach because we lacked complete genome sequences for all the prophage sequences we aimed to analyse. Instead, we used the large and small subunits of the terminase, as recommended by Katoh et al. (2019), as these sequences were the most prevalent across our identified prophage sequences. The integrase was used only for reference genomes and sequences positive for this protein, as recommended by Colavecchio et al. (2017). While other markers like capsid proteins, integrases, and phage spanins have been proposed (Dion et al., 2020; Gorbalenya & Lauber, 2022; Holmes, 2011; Kongari et al., 2018), they were not consistently present in our prophage. Therefore, given the limitations of our data and the complexities of viral phylogenetics, the use of the concatenated terminase subunits represented a pragmatic and justifiable approach to explore the relationships between these prophages. Further research is indeed needed to establish universally accepted methods for viral phylogenetics.

Finally, our research underscores the importance of prophages in *S. enterica* adaptability and their relationship with specific serovars, particularly in poultry farms. Our *in-silico* approach represents a novel way to investigate these prophages and combining this with experimental assays could significantly aid in the development of targeted phage therapies.

## Data availability

*S. enterica* sequenced genomes are avialable under bioproject PRJEB37560 in the NCBI. The corresponding codification and serovar of every *S. enterica* genome and the quality control measures are presented in Supplementary Table 1. The custom database built for this study is listed with their corresponding accession numbers to NCBI in Supplementary Table 2. A list of every prophage detected for every *S. enterica* genome along with their corresponding codification and serovar. All alignments and other raw data used for the analysis in this study are deposited at https://github.com/fran1722/Exploring-prophages-in-Salmonella-enterica-an-in-silico-approach.

## Acknowledgements

Our acknowledgements to Mateo Carvajal and Belén Prado from the Bioinformatics lab at Universidad San Francisco de Quito for their guidance and teachings in basic bioinformatics. We also thank Dr. William Calero at Universidad Técnica de Ambato for his assistance in choosing the proper bioinformatic tools for the analysis of prophage genomes and their genome annotation.

---

**Supplementary Figure 1.**
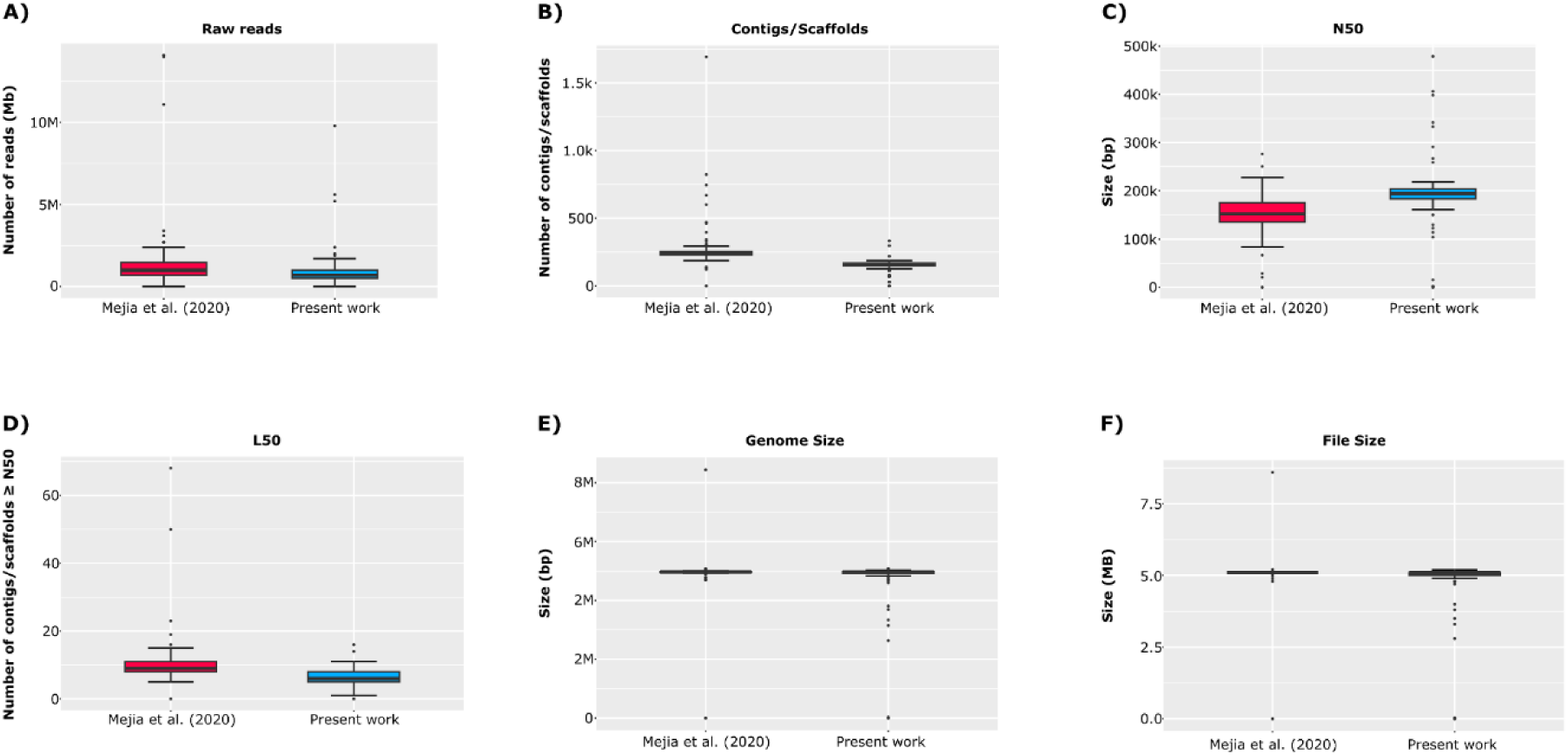
Box plots illustrating the differences between the prior study by Mejia et al. (2020) (depicted in red) and the current work (depicted in blue), focusing on various assembly and quality control statistics. A) Differences in the number of reads obtained after quality control for raw reads. B) Differences in the number of contigs/scaffolds obtained. C) Differences in N50 measured in fragment size. D) Differences in L50 measured in the number of contigs/scaffolds ≥ N50. E) Differences in genome size obtained. F) Differences in file size obtained.

**Supplementary Figure 2.**
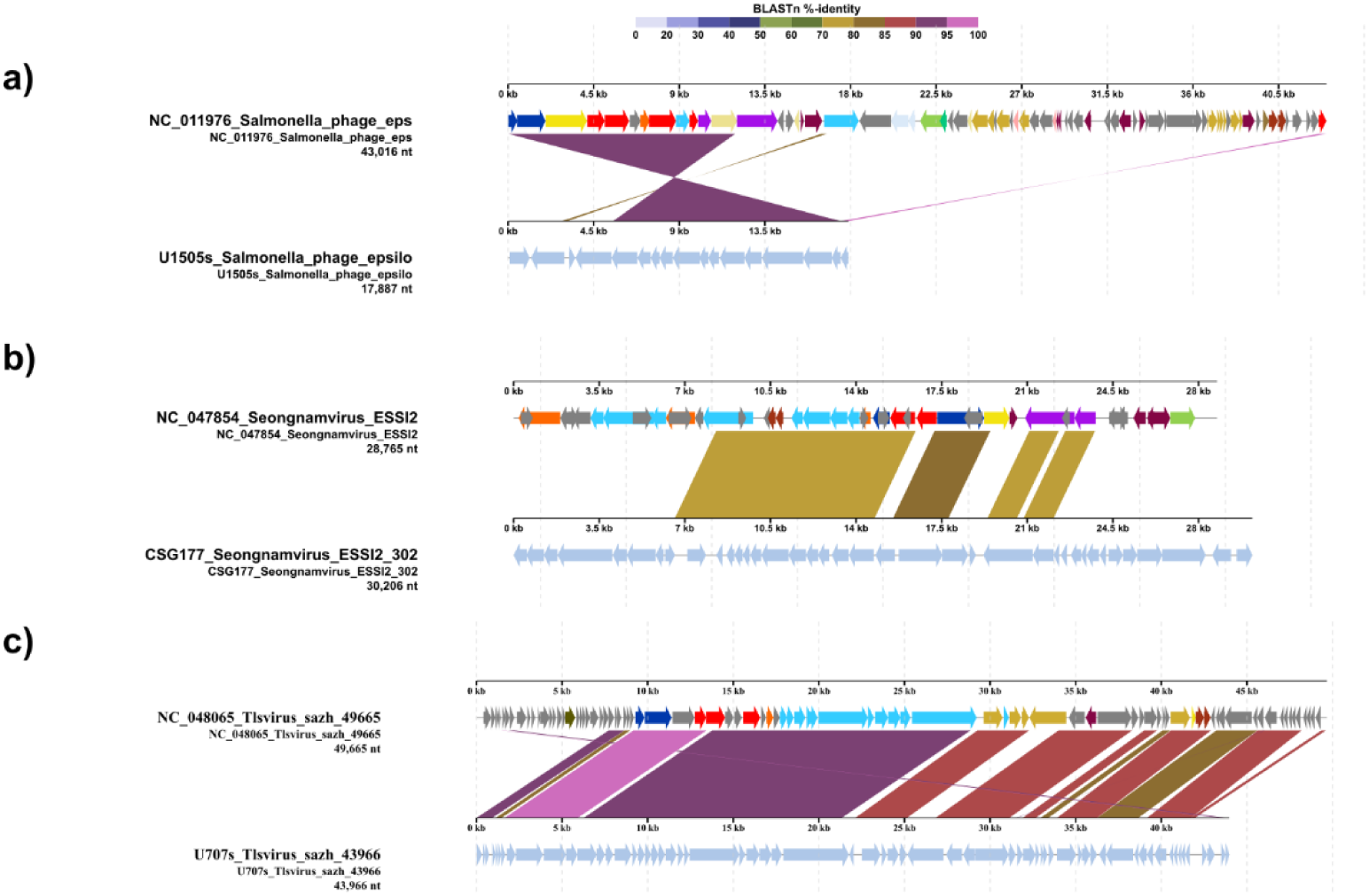
Comparative genomics of a) Salmonella phage epsilon34-like, b) Seongnamvirus ESSI2-like, and c) Tlsvirus sazh-like sequences of varying sizes detected in S. enterica. The legend on the right indicates the main features identified for these prophage sequences compared to the reference genomes. At the top, a BLAST %-identity bar indicates the identity percentage of every genetic block or region compared to the reference genome, according to the color palette on the top. Obtained using DigAlign (v2.0).

**Supplementary Figure 3.**
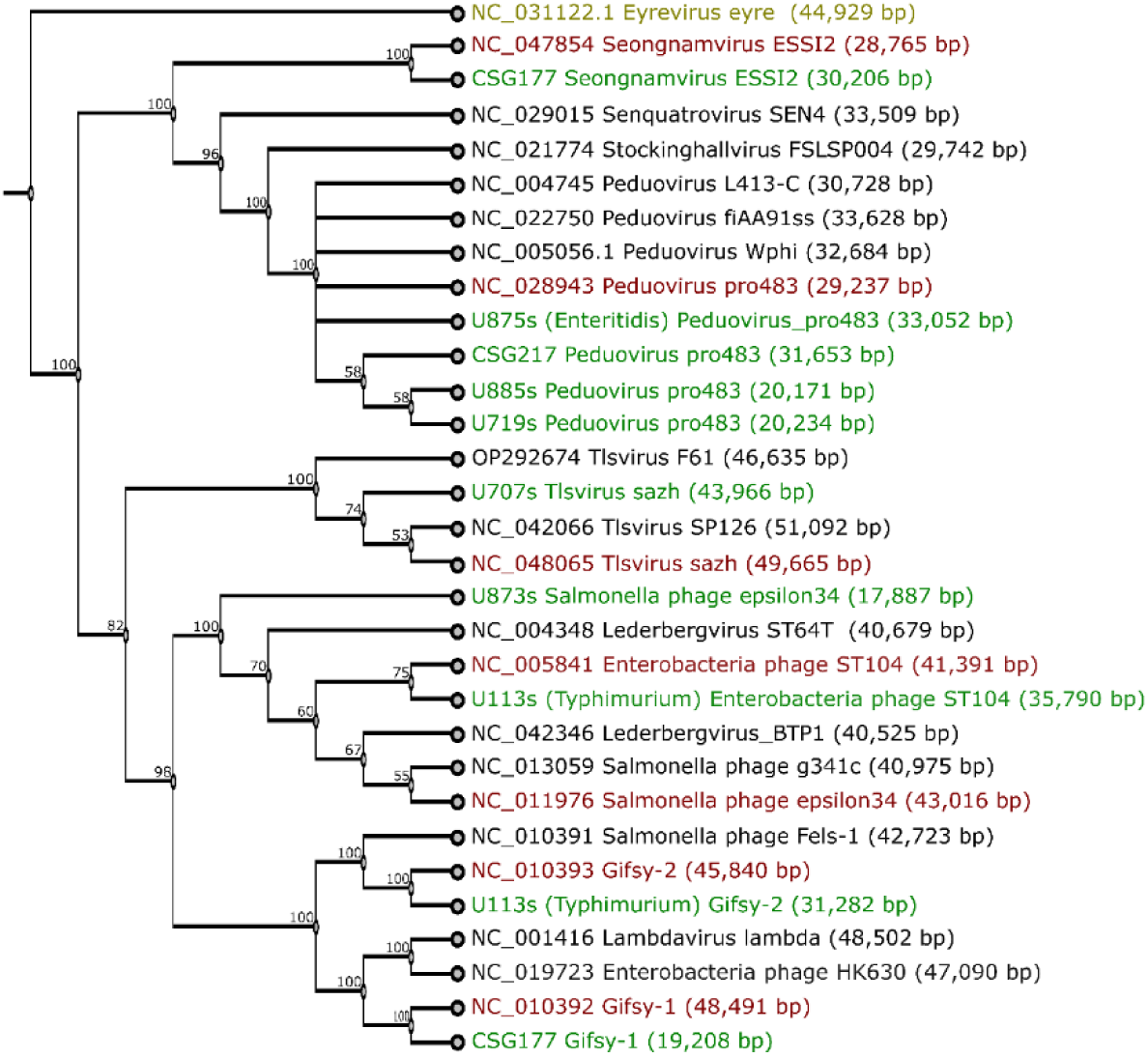
Neighbor-joining tree constructed with terminase small subunit (TerS) amino acid sequences. The outgroup is indicated in yellow, the prophage reference genomes are indicated in red, and the obtained prophage sequences are depicted in green. The TerS sequences were extracted after annotation from every representative prophage sequence detected, their corresponding reference and other related prophage genomes. Obtained using FigTree (v.1.4.4).

**Supplementary Figure 4.**
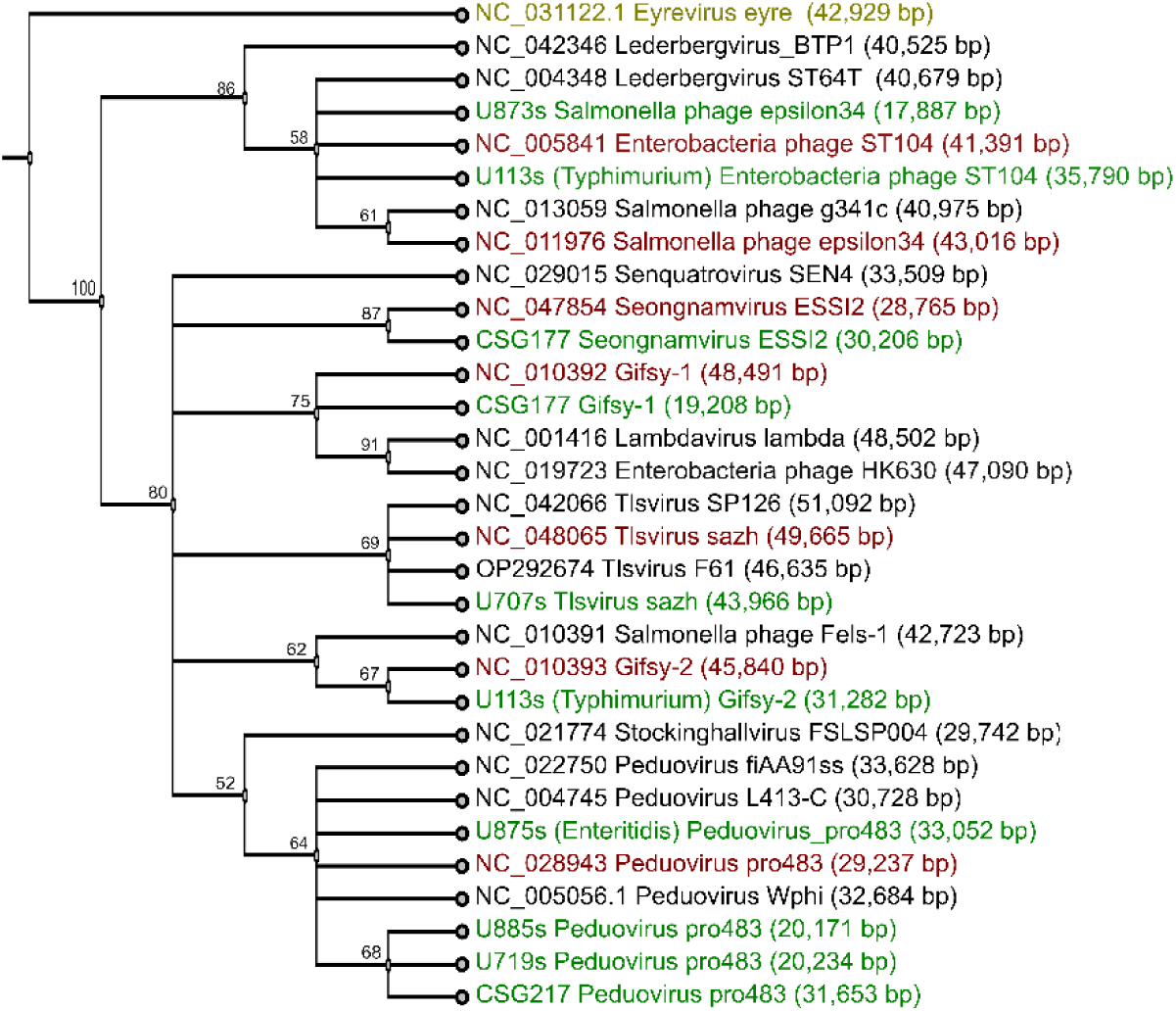
Neighbor-joining tree constructed with terminase large subunit (TerL) amino acid sequences. The outgroup is indicated in yellow, the prophage reference genomes are indicated in red, and the obtained prophage sequences are depicted in green. The TerL sequences were extracted after annotation from every representative prophage sequence detected, their corresponding reference and other related prophage genomes. Obtained using FigTree (v.1.4.4).

**Supplementary Table 1.** Details for all Salmonella enterica genomes assembled, their quality control statistics and indicators.

| Sample name | NCBI Code | Serovar | File Size before trimming (MB) |  | File Size after trimming (MB) |  | Assembly summary: abyss-fac |  |  |  |  |  |  |  |  |  | BUSCO | Assembly: SPAdes |
| --- | --- | --- | --- | --- | --- | --- | --- | --- | --- | --- | --- | --- | --- | --- | --- | --- | --- | --- |
|  |  |  | Forward | Reverse | Forward | Reverse | n | n:500 | L50 | min | N75 | N50 | N25 | E-size | max | sum | Genome Complete (%) | Assembled File Size (MB) |
| Se_Q_001 | CSG217 | *S. Infantis* | 103 | 105 | 73 | 74 | 157 | 62 | 5 | 515 | 138387 | 266844 | 470431 | 451299 | 1195276 | 4982149 | 98,4 | 5,1 |
| Se_Q_002 | CSG232 | *S. Typhimurium* | 92 | 93 | 65 | 66 | 69 | 28 | 4 | 545 | 229450 | 478958 | 647238 | 499866 | 978044 | 4698564 | 98,4 | 4,8 |
| Se_Q_003 | CSG252 | *S. Typhimurium* | 69 | 70 | 49 | 49 | 80 | 33 | 5 | 545 | 171440 | 406336 | 478270 | 427904 | 977752 | 4697863 | 98,4 | 4,8 |
| Se_Q_004 | CSG177 | *S. Infantis* | 58 | 60 | 41 | 42 | 178 | 79 | 8 | 523 | 149779 | 259203 | 340779 | 233524 | 376736 | 4925717 | 98,4 | 5,1 |
| Se_Q_005 | CSG175 | *S. Infantis* | 59 | 60 | 42 | 43 | 151 | 63 | 8 | 501 | 111884 | 194191 | 385887 | 228439 | 525933 | 4706473 | 98,4 | 4,8 |
| Se_Q_006 | CSG132 | *S. Infantis* | 63 | 64 | 44 | 45 | 157 | 72 | 9 | 503 | 111845 | 181160 | 226718 | 253770 | 719507 | 4914638 | 98,4 | 5 |
| Se_Q_007 | U113s | *S. Typhimurium* | 57 | 58 | 40 | 41 | 178 | 89 | 10 | 537 | 83215 | 178330 | 266195 | 179917 | 333658 | 4955233 | 99,2 | 5,1 |
| Se_Q_008 | U114s | *S. Enteritidis* | 56 | 57 | 39 | 40 | 79 | 39 | 6 | 542 | 153089 | 291342 | 375907 | 302264 | 647270 | 4632449 | 98,4 | 4,8 |
| Se_Q_009 | U120s | *S. Infantis* | 116 | 118 | 81 | 82 | 176 | 77 | 8 | 515 | 89905 | 150188 | 474554 | 335336 | 1029493 | 5083069 | 100,0 | 5,2 |
| Se_Q_010 | U121s | *S. Infantis* | 54 | 55 | 38 | 39 | 153 | 78 | 9 | 523 | 103540 | 183601 | 323905 | 222476 | 554236 | 4871727 | 98,4 | 5 |
| Se_Q_011 | U123s | *S. Infantis* | 49 | 50 | 35 | 36 | 137 | 78 | 11 | 523 | 58371 | 129590 | 183859 | 123842 | 225091 | 3686054 | 90,3 | 3,8 |
| Se_Q_012 | U127s | *S. Infantis* | 55 | 56 | 38 | 39 | 143 | 72 | 8 | 617 | 105437 | 192292 | 333656 | 237973 | 617933 | 4782068 | 100,0 | 4,9 |
| Se_Q_013 | U128s | *S. Infantis* | 78 | 80 | 54 | 55 | 163 | 66 | 6 | 515 | 115018 | 201642 | 371030 | 417143 | 1195275 | 4962512 | 98,4 | 5,1 |
| Se_Q_014 | U129s | *S. Infantis* | 94 | 96 | 68 | 69 | 156 | 72 | 8 | 515 | 112205 | 165469 | 438969 | 262897 | 604074 | 5082850 | 100,0 | 5,2 |
| Se_Q_015 | U219s | *S. Infantis* | 76 | 78 | 55 | 55 | 167 | 75 | 9 | 515 | 111806 | 180235 | 324280 | 228602 | 590272 | 5077223 | 100,0 | 5,2 |
| Se_Q_016 | U634s | *S. Infantis* | 49 | 50 | 35 | 36 | 129 | 63 | 10 | 515 | 105545 | 183598 | 237218 | 191325 | 492353 | 4597871 | 96,8 | 4,7 |
| Se_Q_017 | U638s | *S. Infantis* | 73 | 74 | 52 | 53 | 159 | 71 | 6 | 502 | 114987 | 183947 | 525590 | 429418 | 1195182 | 4973292 | 100,0 | 5,1 |
| Se_Q_018 | U639s | *S. Infantis* | 74 | 75 | 53 | 53 | 152 | 71 | 8 | 515 | 114952 | 183598 | 370826 | 250527 | 589934 | 4955057 | 98,4 | 5,1 |
| Se_Q_019 | U652s | *S. Infantis* | 73 | 74 | 52 | 53 | 148 | 67 | 6 | 515 | 103469 | 201608 | 370969 | 411990 | 1195009 | 5001213 | 100,0 | 5,1 |
| Se_Q_021 | U664s | *S. Infantis* | 97 | 98 | 68 | 69 | 147 | 132 | 14 | 542 | 56723 | 113808 | 187395 | 123342 | 313697 | 4887325 | 100,0 | 5 |
| Se_Q_022 | U666s | *S. Infantis* | 62 | 63 | 44 | 44 | 166 | 75 | 8 | 503 | 111806 | 194184 | 324230 | 256584 | 719600 | 4949368 | 100,0 | 5,1 |
| Se_Q_023 | U669s | *S. Infantis* | 64 | 65 | 46 | 46 | 152 | 68 | 6 | 503 | 103482 | 194191 | 525799 | 377336 | 1028952 | 4943934 | 100,0 | 5,1 |
| Se_Q_024 | U672s | *S. Infantis* | 72 | 73 | 51 | 52 | 165 | 71 | 7 | 552 | 94393 | 183600 | 525862 | 278981 | 719785 | 4972344 | 100,0 | 5,1 |
| Se_Q_025 | U676s | *S. Infantis* | 59 | 60 | 42 | 43 | 173 | 69 | 8 | 515 | 105321 | 180609 | 525731 | 288308 | 824615 | 4919924 | 100,0 | 5 |
| Se_Q_026 | U679s | *S. Infantis* | 106 | 108 | 75 | 76 | 149 | 67 | 5 | 515 | 105545 | 183599 | 525922 | 439834 | 1192981 | 4941018 | 99,2 | 5,1 |
| Se_Q_027 | U682s | *S. Infantis* | 88 | 89 | 62 | 62 | 156 | 66 | 6 | 515 | 124631 | 183687 | 469482 | 370959 | 1029473 | 4966517 | 100,0 | 5,1 |
| Se_Q_028 | U684s | *S. Infantis* | 72 | 73 | 51 | 51 | 160 | 73 | 8 | 501 | 93967 | 183687 | 438783 | 252912 | 589935 | 4938821 | 99,2 | 5,1 |
| Se_Q_029 | U689s | *S. Infantis* | 73 | 74 | 52 | 52 | 155 | 64 | 7 | 552 | 111884 | 183947 | 525921 | 362993 | 1029044 | 4963870 | 99,2 | 5,1 |
| Se_Q_030 | U692s | *S. Infantis* | 79 | 81 | 55 | 56 | 162 | 67 | 6 | 512 | 105437 | 194416 | 525918 | 380846 | 1044953 | 4962358 | 98,4 | 5,1 |
| Se_Q_031 | U706s | *S. Infantis* | 65 | 66 | 46 | 46 | 152 | 76 | 9 | 515 | 96311 | 181513 | 324225 | 208948 | 521940 | 4898829 | 100,0 | 5 |
| Se_Q_032 | U707s | *S. Infantis* | 77 | 79 | 55 | 56 | 170 | 75 | 9 | 515 | 93966 | 180521 | 368507 | 214594 | 522039 | 4956781 | 100,0 | 5,1 |
| Se_Q_033 | U708s | *S. Infantis* | 73 | 74 | 51 | 52 | 149 | 60 | 7 | 501 | 111884 | 194278 | 438876 | 273065 | 590544 | 4969235 | 100,0 | 5,1 |
| Se_Q_034 | U711s | *S. Infantis* | 55 | 56 | 39 | 39 | 150 | 77 | 9 | 515 | 105321 | 183859 | 268385 | 206387 | 471316 | 4936538 | 100,0 | 5,1 |
| Se_Q_035 | U712s | *S. Infantis* | 96 | 97 | 68 | 69 | 153 | 68 | 6 | 515 | 96310 | 201656 | 371360 | 414389 | 1195275 | 4971725 | 98,4 | 5,1 |
| Se_Q_036 | U715s | *S. Infantis* | 141 | 144 | 99 | 101 | 165 | 63 | 5 | 515 | 127845 | 203919 | 469779 | 444181 | 1195182 | 4975527 | 99,2 | 5,1 |
| Se_Q_038 | U719s | *S. Infantis* | 198 | 200 | 134 | 136 | 158 | 65 | 6 | 515 | 105545 | 183688 | 444040 | 419315 | 1179805 | 4953029 | 98,4 | 5,1 |
| Se_Q_039 | U723s | *S. Infantis* | 120 | 123 | 78 | 80 | 169 | 70 | 6 | 515 | 105321 | 183687 | 470132 | 371174 | 1029473 | 4966517 | 100,0 | 5,1 |
| Se_Q_041 | U728s | *S. Infantis* | 179 | 182 | 122 | 124 | 168 | 66 | 5 | 507 | 105545 | 203918 | 525853 | 448781 | 1188867 | 4972216 | 100,0 | 5,1 |
| Se_Q_042 | U729s | *S. Infantis* | 105 | 107 | 74 | 75 | 163 | 67 | 6 | 515 | 105321 | 183600 | 525862 | 381778 | 1029474 | 4965114 | 99,2 | 5,1 |
| Se_Q_043 | U758s | *S. Infantis* | 92 | 94 | 64 | 65 | 161 | 71 | 8 | 515 | 103520 | 183600 | 370969 | 254967 | 604576 | 4947710 | 99,2 | 5,1 |
| Se_Q_044 | U759s | *S. Infantis* | 97 | 99 | 67 | 68 | 155 | 63 | 6 | 515 | 148183 | 201638 | 443813 | 432023 | 1195275 | 4973905 | 99,2 | 5,1 |
| Se_Q_045 | U762s | *S. Infantis* | 115 | 117 | 70 | 71 | 156 | 63 | 6 | 515 | 111885 | 203916 | 470180 | 423071 | 1195182 | 4967830 | 99,2 | 5,1 |
| Se_Q_046 | U769s | *S. Infantis* | 94 | 95 | 66 | 67 | 137 | 64 | 6 | 515 | 105438 | 201620 | 444101 | 426266 | 1195275 | 4974672 | 99,2 | 5,1 |
| Se_Q_047 | U773s | *S. Infantis* | 89 | 90 | 63 | 64 | 156 | 67 | 8 | 515 | 93967 | 183599 | 371597 | 253888 | 649759 | 4970622 | 98,4 | 5,1 |
| Se_Q_048 | U801s | *S. Infantis* | 161 | 164 | 110 | 111 | 152 | 61 | 5 | 515 | 114988 | 203915 | 525921 | 451228 | 1187838 | 4956900 | 98,4 | 5,1 |
| Se_Q_049 | U804s | *S. Infantis* | 142 | 144 | 99 | 101 | 171 | 67 | 5 | 504 | 105545 | 203915 | 470256 | 446519 | 1195089 | 4942619 | 98,4 | 5,1 |
| Se_Q_050 | U806s | *S. Infantis* | 101 | 102 | 71 | 72 | 161 | 62 | 6 | 504 | 148183 | 201623 | 444039 | 435349 | 1200177 | 4970943 | 100,0 | 5,1 |
| Se_Q_051 | U814s | *S. Infantis* | 192 | 195 | 131 | 133 | 147 | 65 | 5 | 515 | 105429 | 203915 | 525858 | 446007 | 1195273 | 4973872 | 98,4 | 5,1 |
| Se_Q_052 | U815s | *S. Infantis* | 95 | 96 | 66 | 67 | 177 | 65 | 6 | 515 | 93904 | 183597 | 470263 | 370584 | 1044861 | 4925562 | 99,2 | 5,1 |
| Se_Q_053 | U818s | *S. Infantis* | 114 | 116 | 81 | 82 | 168 | 66 | 6 | 515 | 89905 | 201627 | 444040 | 417117 | 1195271 | 4974881 | 99,2 | 5,1 |
| Se_Q_054 | U855s | *S. Infantis* | 104 | 106 | 49 | 49 | 146 | 65 | 7 | 515 | 105321 | 183687 | 525901 | 361229 | 1029084 | 4931517 | 99,2 | 5,1 |
| Se_Q_055 | U857s | *S. Infantis* | 101 | 103 | 71 | 72 | 159 | 66 | 6 | 515 | 105545 | 201626 | 371299 | 417759 | 1195089 | 4971699 | 100,0 | 5,1 |
| Se_Q_056 | U860s | *S. Infantis* | 75 | 76 | 53 | 54 | 137 | 63 | 7 | 515 | 124631 | 203916 | 371597 | 284253 | 676384 | 4916076 | 98,4 | 5 |
| Se_Q_057 | U863s | *S. Infantis* | 1,2 GB | 1,2 GB | 820 | 829 | 153 | 60 | 5 | 515 | 127741 | 203920 | 525828 | 454696 | 1195275 | 4971725 | 100,0 | 5,1 |
| Se_Q_058 | U865s | *S. Infantis* | 148 | 151 | 105 | 106 | 166 | 69 | 6 | 515 | 92521 | 183687 | 470134 | 395552 | 1099117 | 4973961 | 99,2 | 5,1 |
| Se_Q_059 | U866s | *S. Infantis* | 75 | 76 | 53 | 54 | 156 | 70 | 6 | 502 | 103519 | 201621 | 444101 | 423020 | 1195097 | 4976315 | 98,4 | 5,1 |
| Se_Q_061 | U871s | *S. Infantis* | 71 | 72 | 51 | 51 | 183 | 76 | 8 | 502 | 103515 | 183859 | 371030 | 311801 | 942896 | 4959522 | 100,0 | 5,1 |
| Se_Q_062 | U873s | *S. Infantis* | 112 | 114 | 78 | 80 | 163 | 67 | 6 | 515 | 127731 | 201644 | 444101 | 420547 | 1191517 | 4976344 | 97,6 | 5,1 |
| Se_Q_063 | U875s | *S. Enteritidis* | 147 | 150 | 104 | 106 | 76 | 30 | 3 | 618 | 185781 | 406400 | 1455556 | 672327 | 1455556 | 4773271 | 98,4 | 4,9 |
| Se_Q_064 | U878s | *S. Infantis* | 82 | 84 | 58 | 58 | 169 | 69 | 7 | 515 | 96311 | 181222 | 525954 | 358858 | 1029772 | 4977338 | 99,2 | 5,1 |
| Se_Q_065 | U880s | *S. Infantis* | 104 | 106 | 75 | 76 | 160 | 63 | 5 | 515 | 105545 | 203916 | 525868 | 451211 | 1195275 | 5016594 | 100,0 | 5,1 |
| Se_Q_066 | U883s | *S. Infantis* | 80 | 82 | 57 | 58 | 156 | 63 | 5 | 515 | 114968 | 203919 | 525791 | 444421 | 1194908 | 4971129 | 99,2 | 5,1 |
| Se_Q_067 | U885s | *S. Infantis* | 78 | 80 | 55 | 56 | 141 | 68 | 7 | 515 | 105545 | 201635 | 370969 | 328926 | 960342 | 4961154 | 100,0 | 5,1 |
| Se_Q_068 | U888s | *S. Infantis* | 204 | 207 | 145 | 147 | 167 | 67 | 6 | 515 | 92521 | 183600 | 470196 | 418552 | 1145098 | 4973494 | 98,4 | 5,1 |
| Se_Q_069 | U889s | *S. Infantis* | 59 | 60 | 42 | 42 | 149 | 63 | 10 | 515 | 114982 | 183687 | 226203 | 202617 | 510646 | 4723271 | 98,4 | 4,9 |
| Se_Q_070 | U890s | *S. Infantis* | 79 | 81 | 56 | 57 | 170 | 69 | 8 | 515 | 113153 | 183687 | 371360 | 300266 | 915601 | 4972832 | 100,0 | 5,1 |
| Se_Q_071 | U895s | S. Infantis | 61 | 62 | 43 | 44 | 142 | 66 | 7 | 515 | 124631 | 203915 | 525920 | 298640 | 743739 | 4928924 | 100,0 | 5 |
| Se_Q_072 | U897s | S. Infantis | 56 | 57 | 40 | 40 | 147 | 65 | 8 | 515 | 111884 | 194406 | 438876 | 239230 | 510645 | 4869524 | 100,0 | 5 |
| Se_Q_074 | U969s | S. Infantis | 70 | 71 | 49 | 49 | 160 | 68 | 8 | 515 | 103470 | 183687 | 277902 | 270230 | 807038 | 4927197 | 99,2 | 5 |
| Se_Q_075 | U972s | S. Infantis | 88 | 90 | 63 | 64 | 159 | 66 | 7 | 504 | 92095 | 183600 | 371658 | 345249 | 1029138 | 4972348 | 98,4 | 5,1 |
| Se_Q_076 | U973s | S. Infantis | 66 | 67 | 33 | 33 | 112 | 52 | 7 | 523 | 89700 | 183599 | 224894 | 164281 | 298879 | 3144061 | 87,1 | 3,3 |
| Se_Q_078 | U979s | S. Infantis | 73 | 74 | 52 | 53 | 148 | 61 | 6 | 502 | 127843 | 203917 | 525795 | 372931 | 1028771 | 4830648 | 98,4 | 5 |
| Se_Q_079 | U980s | S. Infantis | 72 | 73 | 51 | 51 | 145 | 71 | 8 | 515 | 114902 | 183687 | 525908 | 271306 | 719606 | 4968981 | 99,2 | 5,1 |
| Se_Q_080 | U983s | S. Infantis | 87 | 88 | 62 | 63 | 158 | 68 | 7 | 503 | 115007 | 183860 | 438969 | 256380 | 604576 | 4950623 | 99,2 | 5,1 |
| Se_Q_081 | U986s | S. Infantis | 89 | 90 | 63 | 64 | 166 | 69 | 7 | 515 | 145663 | 194156 | 444101 | 282854 | 590383 | 4971259 | 100,0 | 5,1 |
| Se_Q_082 | U998s | S. Infantis | 58 | 59 | 41 | 41 | 169 | 74 | 7 | 515 | 103500 | 180946 | 470149 | 319161 | 897909 | 4930452 | 99,2 | 5,1 |
| Se_Q_083 | U1000s | S. Infantis | 151 | 154 | 107 | 109 | 170 | 65 | 5 | 515 | 96310 | 203916 | 470191 | 434277 | 1177679 | 4958205 | 98,4 | 5,1 |
| Se_Q_084 | U1003s | S. Infantis | 65 | 66 | 46 | 47 | 169 | 68 | 6 | 515 | 145663 | 194285 | 525919 | 349482 | 949950 | 4962491 | 99,2 | 5,1 |
| Se_Q_085 | U1006s | S. Infantis | 60 | 61 | 43 | 43 | 170 | 78 | 9 | 503 | 114985 | 180829 | 324316 | 215536 | 510820 | 4928362 | 99,2 | 5,1 |
| Se_Q_086 | U1010s | S. Infantis | 217 | 222 | 147 | 150 | 157 | 63 | 5 | 515 | 127966 | 203919 | 525767 | 453608 | 1195274 | 4970923 | 100,0 | 5,1 |
| Se_Q_087 | U1019s | S. Infantis | 175 | 180 | 121 | 123 | 152 | 60 | 5 | 515 | 127835 | 203831 | 470164 | 440745 | 1195276 | 5003523 | 100,0 | 5,1 |
| Se_Q_088 | U1051s | S. Infantis | 142 | 145 | 96 | 98 | 157 | 58 | 5 | 515 | 105429 | 203919 | 525920 | 452315 | 1193237 | 4976698 | 98,4 | 5,1 |
| Se_Q_089 | U1052s | S. Infantis | 147 | 149 | 100 | 101 | 153 | 55 | 5 | 515 | 114978 | 218061 | 525850 | 459568 | 1195276 | 4910636 | 98,4 | 5 |
| Se_Q_090 | U1056s | S. Infantis | 138 | 141 | 96 | 98 | 155 | 64 | 5 | 515 | 105545 | 203916 | 470195 | 438765 | 1195275 | 4974139 | 100,0 | 5,1 |
| Se_Q_091 | U1057s | S. Infantis | 1,3 GB | 1,3 GB | 444 | 452 | 157 | 57 | 5 | 515 | 127741 | 203922 | 525792 | 455204 | 1195276 | 4958329 | 98,4 | 5,1 |
| Se_Q_092 | U1059s | S. Infantis | 186 | 190 | 129 | 131 | 156 | 63 | 6 | 515 | 115806 | 183687 | 469954 | 428704 | 1195269 | 4979617 | 98,4 | 5,1 |
| Se_Q_093 | U1066s | S. Infantis | 235 | 239 | 157 | 160 | 175 | 67 | 7 | 515 | 115806 | 183687 | 328299 | 411906 | 1195275 | 4978508 | 98,4 | 5,1 |
| Se_Q_094 | U1068s | S. Infantis | 139 | 141 | 91 | 92 | 185 | 75 | 8 | 515 | 75953 | 194484 | 371300 | 236424 | 605857 | 4969036 | 99,2 | 5,1 |
| Se_Q_095 | U1071s | S. Infantis | 246 | 252 | 170 | 173 | 160 | 66 | 6 | 515 | 105653 | 181631 | 470589 | 439083 | 1195276 | 4973584 | 98,4 | 5,1 |
| Se_Q_096 | U1092s | S. Infantis | 281 | 287 | 74 | 76 | 173 | 76 | 7 | 515 | 92032 | 183599 | 371360 | 362990 | 1096372 | 4945769 | 98,4 | 5,1 |
| Se_Q_097 | U1095s | S. Infantis | 301 | 308 | 208 | 212 | 168 | 61 | 6 | 504 | 105653 | 217634 | 444101 | 426654 | 1195274 | 4942761 | 98,4 | 5,1 |
| Se_Q_098 | U1111s | S. Infantis | 169 | 172 | 119 | 121 | 171 | 62 | 5 | 515 | 105545 | 217656 | 470238 | 454390 | 1200266 | 4983958 | 100,0 | 5,1 |
| Se_Q_099 | U1114s | S. Infantis | 206 | 210 | 146 | 148 | 145 | 59 | 4 | 515 | 116776 | 332783 | 525930 | 463678 | 1195276 | 4970958 | 98,4 | 5,1 |
| Se_Q_100 | U1119s | S. Infantis | 95 | 97 | 65 | 66 | 161 | 66 | 7 | 515 | 105429 | 183687 | 333629 | 409911 | 1195079 | 4958535 | 99,2 | 5,1 |
| Se_Q_101 | U1121s | S. Infantis | 153 | 155 | 108 | 110 | 174 | 64 | 5 | 515 | 96310 | 203919 | 470590 | 441553 | 1195275 | 4975149 | 99,2 | 5,1 |
| Se_Q_102 | U1125s | S. Infantis | 173 | 176 | 124 | 126 | 173 | 65 | 6 | 515 | 105429 | 183647 | 470176 | 431332 | 1195275 | 4971014 | 99,2 | 5,1 |
| Se_Q_103 | U1128s | S. Infantis | 134 | 137 | 95 | 96 | 133 | 49 | 5 | 515 | 127734 | 217930 | 525798 | 462431 | 1195275 | 4918610 | 98,4 | 5 |
| Se_Q_104 | U1132s | S. Infantis | 135 | 137 | 95 | 96 | 182 | 64 | 6 | 515 | 92095 | 183687 | 525921 | 427536 | 1195272 | 4964167 | 98,4 | 5,1 |
| Se_Q_105 | U1133s | S. Infantis | 147 | 150 | 103 | 105 | 166 | 64 | 5 | 515 | 94331 | 203916 | 525876 | 439913 | 1200267 | 5017389 | 100,0 | 5,1 |
| Se_Q_106 | U1135s | S. Infantis | 1010 | 1,1 GB | 480 | 487 | 142 | 59 | 5 | 515 | 145663 | 203922 | 525921 | 458314 | 1195275 | 4972749 | 99,2 | 5,1 |
| Se_Q_107 | U1143s | S. Infantis | 91 | 92 | 64 | 65 | 143 | 63 | 5 | 515 | 124634 | 203920 | 526047 | 451960 | 1190461 | 4973014 | 99,2 | 5,1 |
| Se_Q_108 | U1145s | S. Infantis | 86 | 87 | 61 | 62 | 160 | 65 | 6 | 502 | 105429 | 183687 | 525971 | 371372 | 1029604 | 5001339 | 100,0 | 5,1 |
| Se_Q_109 | U1175s | *S. Infantis* | 66 | 67 | 47 | 47 | 150 | 72 | 9 | 502 | 115034 | 201639 | 280583 | 207648 | 438873 | 4953178 | 98,4 | 5,1 |
| Se_Q_110 | U1178s | *S. Infantis* | 151 | 154 | 105 | 107 | 152 | 70 | 6 | 515 | 105437 | 203888 | 525922 | 327898 | 824584 | 4973125 | 98,4 | 5,1 |
| Se_Q_111 | U1181s | *S. Infantis* | 86 | 89 | 60 | 61 | 133 | 57 | 7 | 515 | 148183 | 201627 | 370742 | 357697 | 1028877 | 4916214 | 98,4 | 5 |
| Se_Q_112 | U1187s | *S. Infantis* | 72 | 74 | 50 | 51 | 174 | 72 | 8 | 515 | 103538 | 180947 | 525800 | 328857 | 951266 | 5012740 | 99,2 | 5,1 |
| Se_Q_113 | U1192s | *S. Infantis* | 73 | 74 | 51 | 52 | 156 | 68 | 7 | 515 | 93967 | 183687 | 438876 | 266279 | 590274 | 4955045 | 98,4 | 5,1 |
| Se_Q_114 | U1193s | *S. Enteritidis* | 88 | 90 | 63 | 63 | 73 | 26 | 4 | 618 | 400868 | 478531 | 694896 | 528125 | 978047 | 4698575 | 98,4 | 4,8 |
| Se_Q_115 | U1196s | *S. Infantis* | 134 | 136 | 93 | 94 | 168 | 64 | 6 | 515 | 115007 | 183687 | 470195 | 433446 | 1195276 | 4971350 | 98,4 | 5,1 |
| Se_Q_116 | U1401 | *S. Infantis* | 67 | 69 | 48 | 48 | 163 | 70 | 8 | 515 | 105429 | 183687 | 269563 | 260207 | 753860 | 4969119 | 100,0 | 5,1 |
| Se_Q_117 | U1405s | *S. Infantis* | 89 | 92 | 62 | 63 | 148 | 63 | 6 | 502 | 148183 | 201620 | 444101 | 430796 | 1195276 | 4973634 | 98,4 | 5,1 |
| Se_Q_118 | U1407s | *S. Infantis* | 141 | 143 | 98 | 99 | 172 | 75 | 6 | 504 | 105429 | 201632 | 444101 | 422133 | 1195182 | 4981059 | 100,0 | 5,1 |
| Se_Q_119 | U1410s | *S. Infantis* | 56 | 58 | 39 | 40 | 138 | 56 | 8 | 515 | 137723 | 194184 | 324237 | 227470 | 443813 | 4662115 | 99,2 | 4,8 |
| Se_Q_120 | U1411s | *S. Infantis* | 66 | 67 | 46 | 47 | 151 | 65 | 8 | 586 | 138005 | 194184 | 383560 | 229155 | 444101 | 4831418 | 98,4 | 5 |
| Se_Q_121 | U1412s | *S. Infantis* | 59 | 61 | 42 | 43 | 333 | 133 | 16 | 508 | 59252 | 103944 | 171465 | 112579 | 224615 | 4710491 | 100,0 | 4,9 |
| Se_Q_122 | U1418s | *S. Infantis* | 98 | 100 | 68 | 69 | 170 | 78 | 6 | 515 | 105545 | 201629 | 371360 | 414892 | 1195276 | 4988717 | 98,4 | 5,1 |
| Se_Q_123 | U1424s | *S. Infantis* | 79 | 81 | 55 | 56 | 172 | 75 | 8 | 515 | 93976 | 181340 | 324484 | 321795 | 1004693 | 4943578 | 100,0 | 5,1 |
| Se_Q_124 | U1431s | *S. Infantis* | 59 | 60 | 41 | 42 | 150 | 71 | 10 | 515 | 114985 | 201626 | 239404 | 198987 | 439942 | 4880964 | 100,0 | 5 |
| Se_Q_125 | U1432s | *S. Infantis* | 57 | 58 | 40 | 40 | 150 | 73 | 9 | 586 | 105321 | 194285 | 232171 | 206695 | 511727 | 4858079 | 98,4 | 5 |
| Se_Q_126 | U1436s | *S. Infantis* | 57 | 58 | 40 | 41 | 127 | 67 | 8 | 515 | 105321 | 194175 | 444101 | 240905 | 521906 | 4872597 | 100,0 | 5 |
| Se_Q_127 | U1437s | *S. Infantis* | 85 | 87 | 59 | 60 | 144 | 64 | 6 | 515 | 145663 | 194191 | 525799 | 380438 | 1029076 | 4968250 | 98,4 | 5,1 |
| Se_Q_128 | U1447s | *S. Infantis* | 85 | 87 | 60 | 61 | 152 | 58 | 7 | 515 | 138104 | 194184 | 438969 | 274360 | 590383 | 4912998 | 98,4 | 5 |
| Se_Q_129 | U1459s | *S. Infantis* | 92 | 94 | 64 | 65 | 180 | 80 | 8 | 515 | 103482 | 180526 | 324266 | 334060 | 1045248 | 4979507 | 100,0 | 5,1 |
| Se_Q_130 | U1467s | *S. Infantis* | 53 | 54 | 37 | 38 | 219 | 93 | 9 | 524 | 47492 | 122678 | 209274 | 155807 | 437028 | 3334242 | 82,3 | 3,5 |
| Se_Q_131 | U1470s | *S. Infantis* | 58 | 60 | 40 | 41 | 147 | 72 | 8 | 523 | 105429 | 194184 | 278475 | 253896 | 653954 | 4911304 | 100,0 | 5 |
| Se_Q_132 | U1473s | *S. Infantis* | 47 | 48 | 33 | 33 | 134 | 56 | 8 | 552 | 87822 | 160941 | 225182 | 199285 | 505523 | 3804469 | 91,9 | 4 |
| Se_Q_133 | U1483s | *S. Infantis* | 98 | 100 | 68 | 70 | 176 | 75 | 7 | 515 | 124227 | 201649 | 438969 | 290447 | 756072 | 4975275 | 98,4 | 5,1 |
| Se_Q_135 | U1493s | *S. Infantis* | 71 | 72 | 49 | 50 | 168 | 76 | 11 | 515 | 82933 | 149868 | 270578 | 211586 | 669891 | 4996844 | 99,2 | 5,1 |
| Se_Q_136 | U1495s | *S. Infantis* | 84 | 85 | 58 | 59 | 179 | 73 | 8 | 515 | 90033 | 166870 | 324680 | 326000 | 1029381 | 4971687 | 98,4 | 5,1 |
| Se_Q_137 | U1496s | *S. Infantis* | 59 | 60 | 42 | 42 | 67 | 31 | 4 | 564 | 225385 | 341534 | 1266045 | 541359 | 1266045 | 4989379 | 99,2 | 5,1 |
| Se_Q_138 | U1505s | *S. Infantis* | 88 | 90 | 63 | 64 | 163 | 76 | 7 | 515 | 107976 | 183687 | 336225 | 340933 | 1029044 | 4946285 | 100,0 | 5,1 |
| Se_Q_139 | U1506s | *S. Infantis* | 61 | 63 | 42 | 43 | 147 | 68 | 8 | 515 | 145663 | 192205 | 324412 | 302782 | 915788 | 4906403 | 99,2 | 5 |
| Se_Q_140 | U1508s | *S. Infantis* | 145 | 148 | 102 | 104 | 138 | 60 | 5 | 515 | 105545 | 203917 | 525920 | 452305 | 1195275 | 4975692 | 99,2 | 5,1 |
| Se_Q_141 | U1519s | *S. Infantis* | 244 | 248 | 171 | 173 | 297 | 109 | 8 | 509 | 105653 | 204075 | 375192 | 234190 | 490266 | 4996052 | 99,2 | 5,1 |
| Se_Q_143 | U1671s | *S. Infantis* | 137 | 139 | 96 | 98 | 160 | 67 | 5 | 515 | 145663 | 203920 | 525920 | 454465 | 1195089 | 4974094 | 100,0 | 5,1 |
| Se_Q_144 | U1672s | *S. Infantis* | 123 | 125 | 87 | 88 | 178 | 69 | 5 | 515 | 105545 | 203916 | 525896 | 441228 | 1195182 | 4979276 | 100,0 | 5,1 |
| Se_Q_145 | U1673s | *S. Infantis* | 80 | 81 | 56 | 57 | 166 | 67 | 7 | 502 | 114905 | 183947 | 443813 | 361724 | 1044874 | 5018874 | 100,0 | 5,1 |
| Se_Q_146 | U1678s | *S. Infantis* | 89 | 91 | 64 | 64 | 161 | 62 | 7 | 502 | 127731 | 183687 | 525737 | 294937 | 719599 | 4962469 | 98,4 | 5,1 |
| Se_Q_147 | U1680s | S. Infantis | 89 | 90 | 61 | 62 | 148 | 53 | 6 | 515 | 124631 | 194757 | 525863 | 389544 | 1029137 | 4915668 | 99,2 | 5 |
| Se_Q_149 | U1683s | S. Infantis | 120 | 122 | 85 | 86 | 152 | 62 | 6 | 515 | 145663 | 201624 | 444101 | 429267 | 1195100 | 4965818 | 99,2 | 5,1 |
| Se_Q_150 | U1686s | S. Infantis | 141 | 143 | 100 | 101 | 163 | 62 | 4 | 515 | 152221 | 398405 | 525799 | 480844 | 1195275 | 5024068 | 99,2 | 5,1 |
| Se_Q_151 | U1688s | S. Infantis | 125 | 127 | 88 | 89 | 187 | 67 | 6 | 515 | 105545 | 183600 | 525879 | 379183 | 1027435 | 4973991 | 99,2 | 5,1 |

**Supplementary Table 2.**
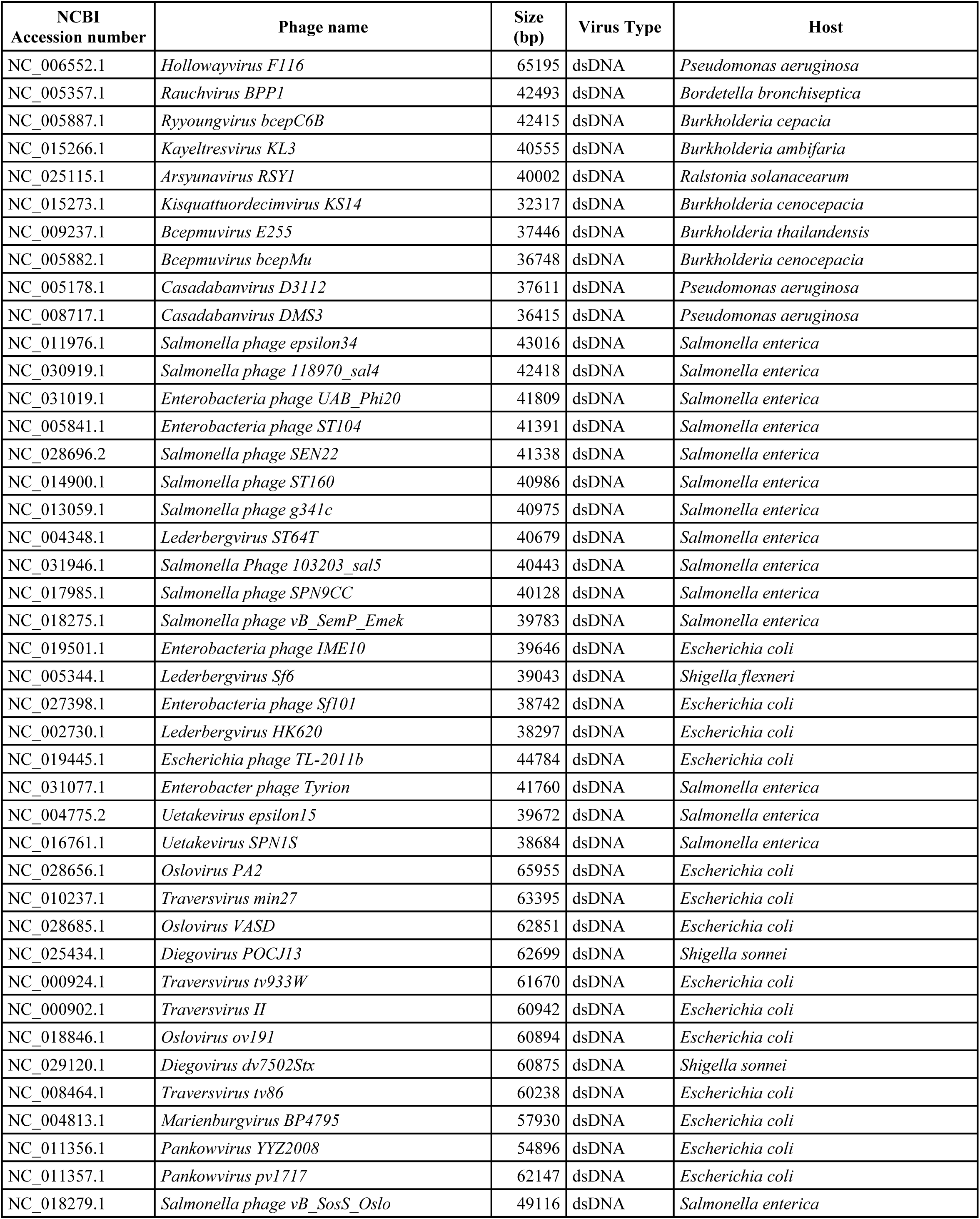
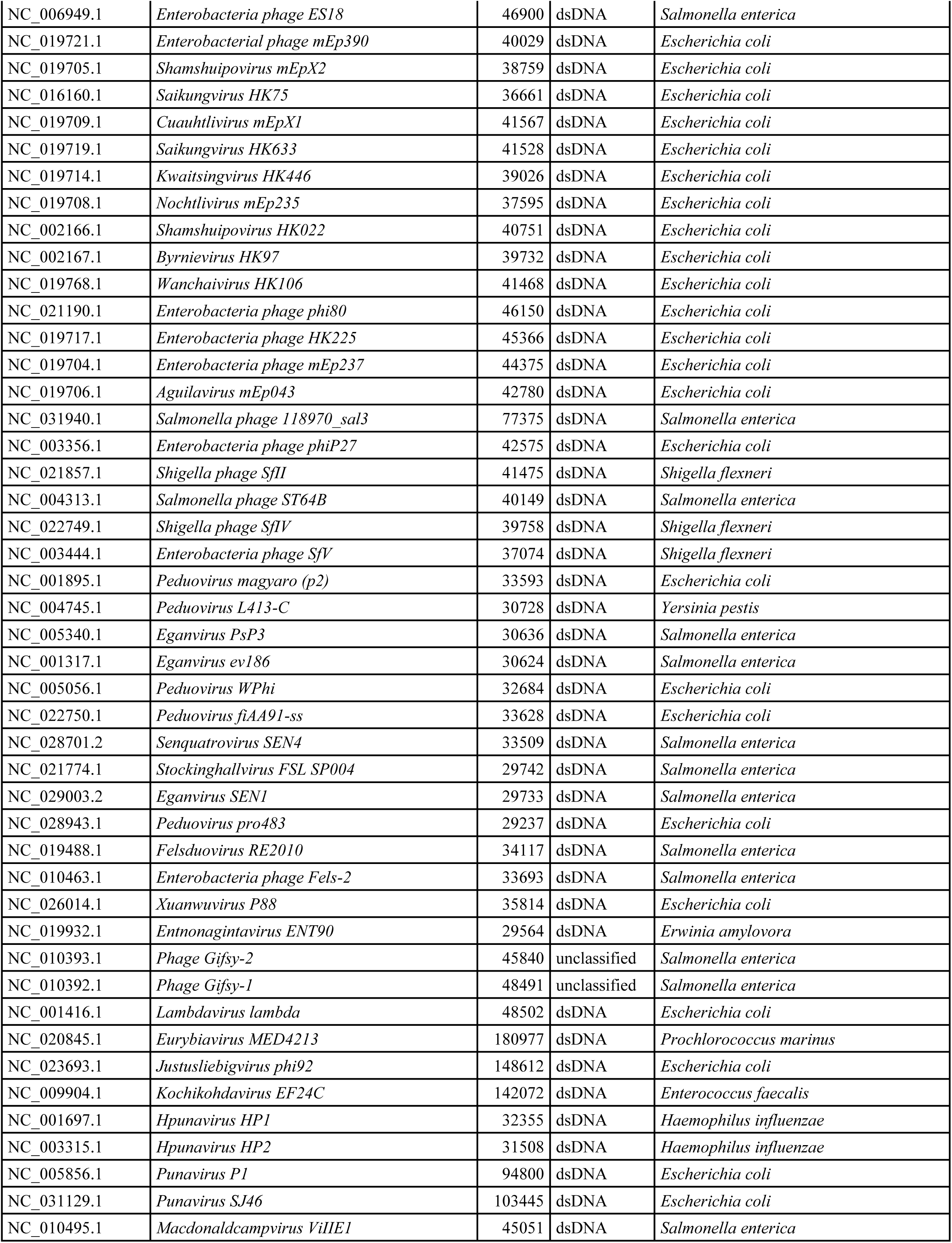
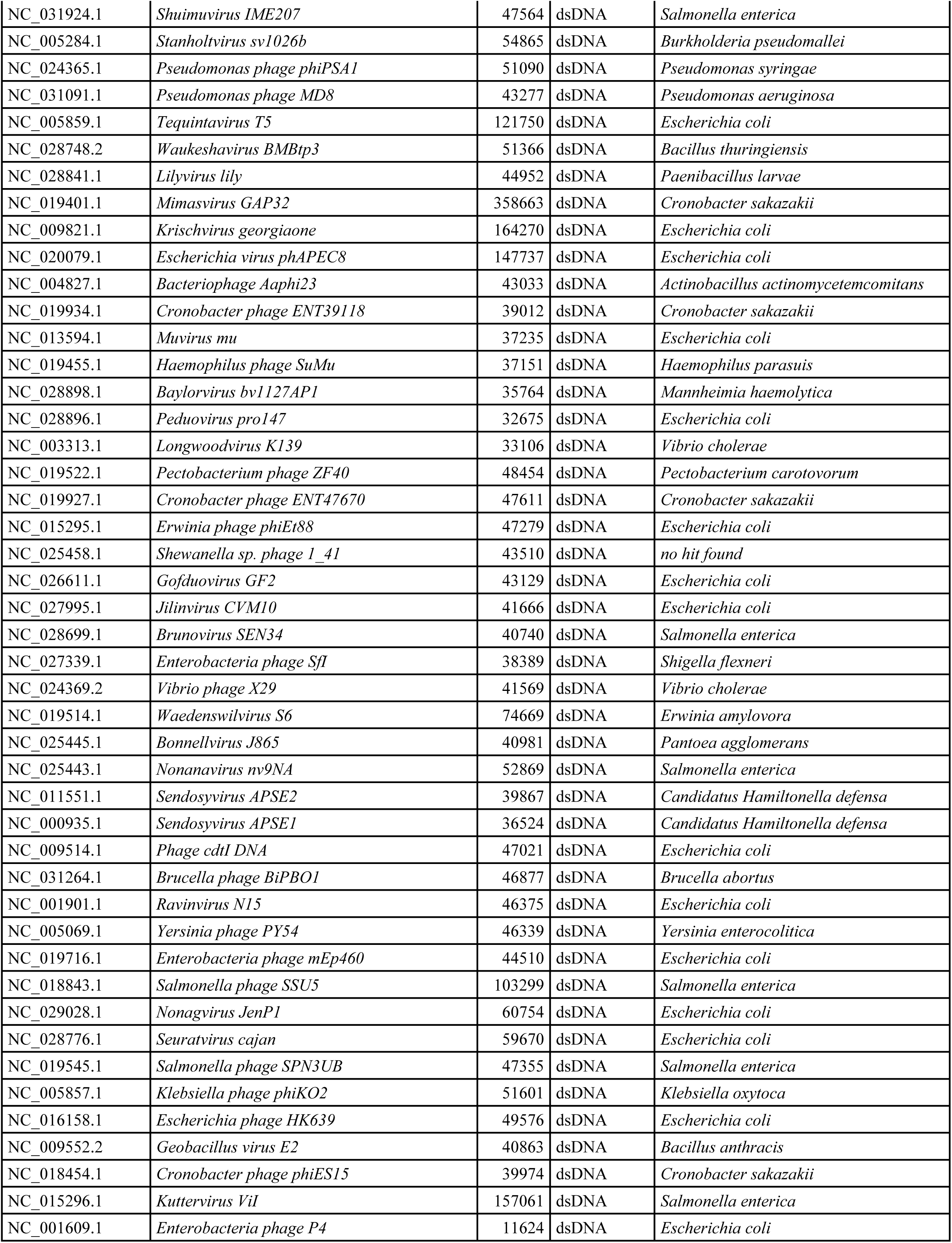
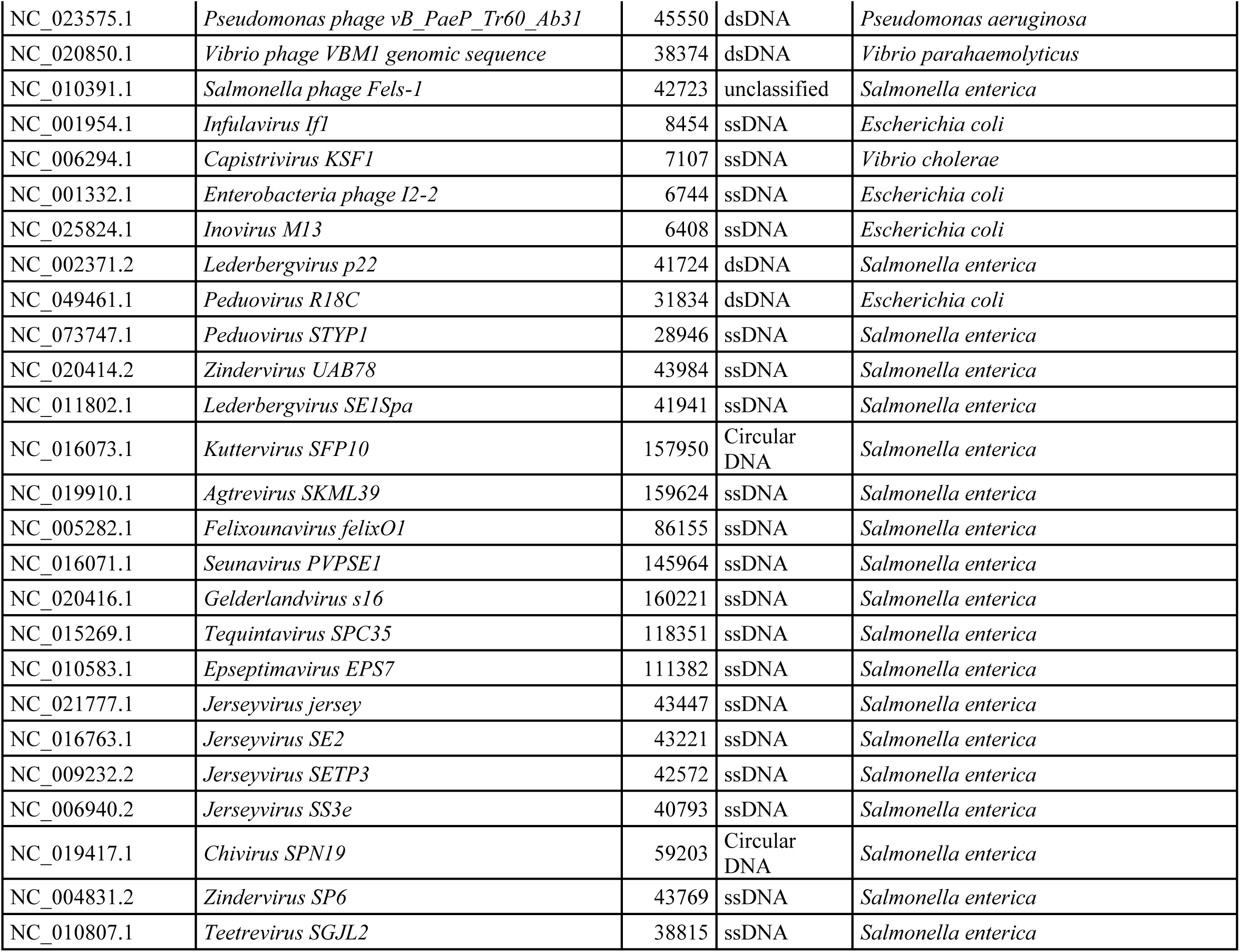
Details of the 160 prophage genomes used for the custom database with information from Switt et al. (2015) and Gao et al. (2020).

**Supplementary Table 3.**
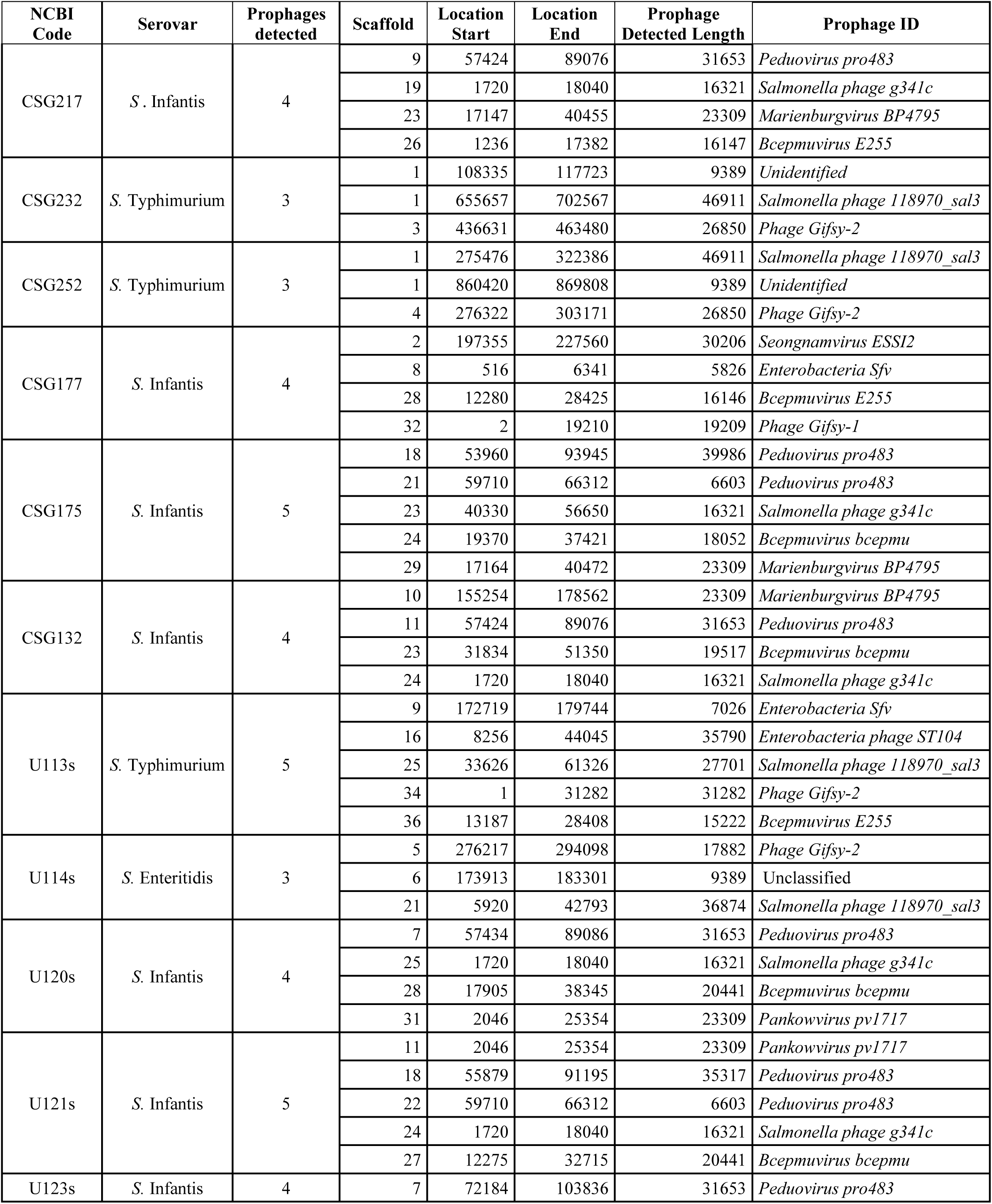
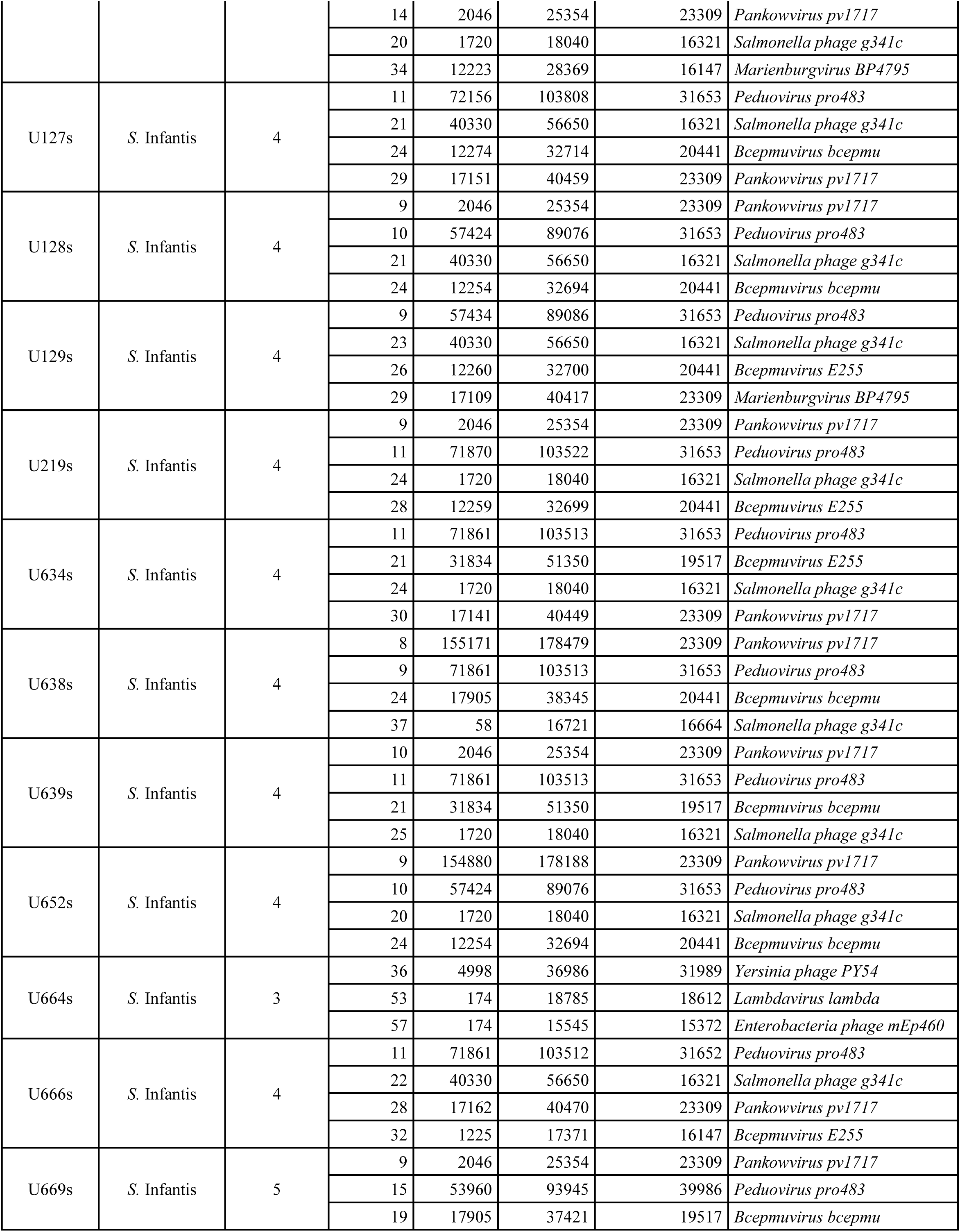
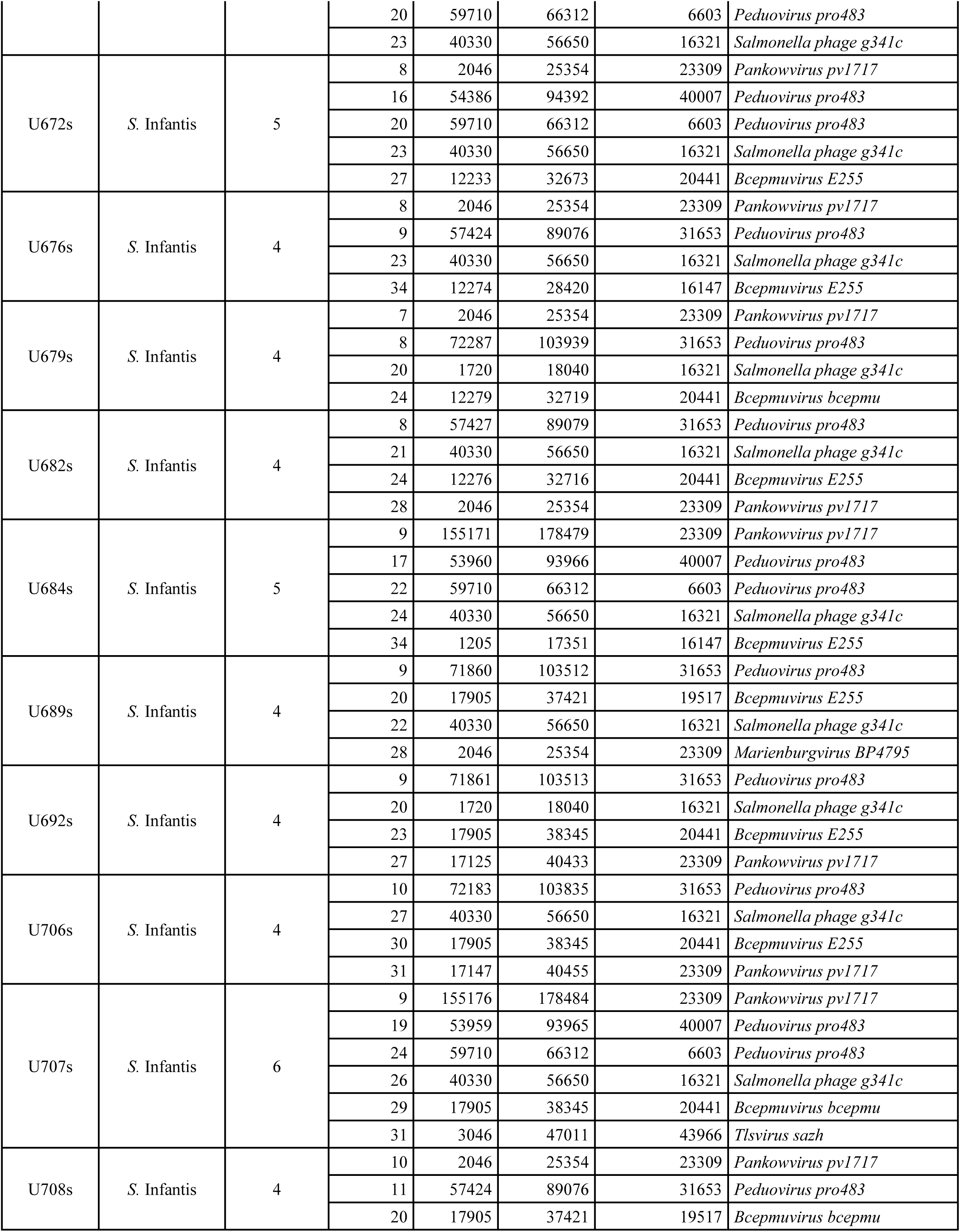
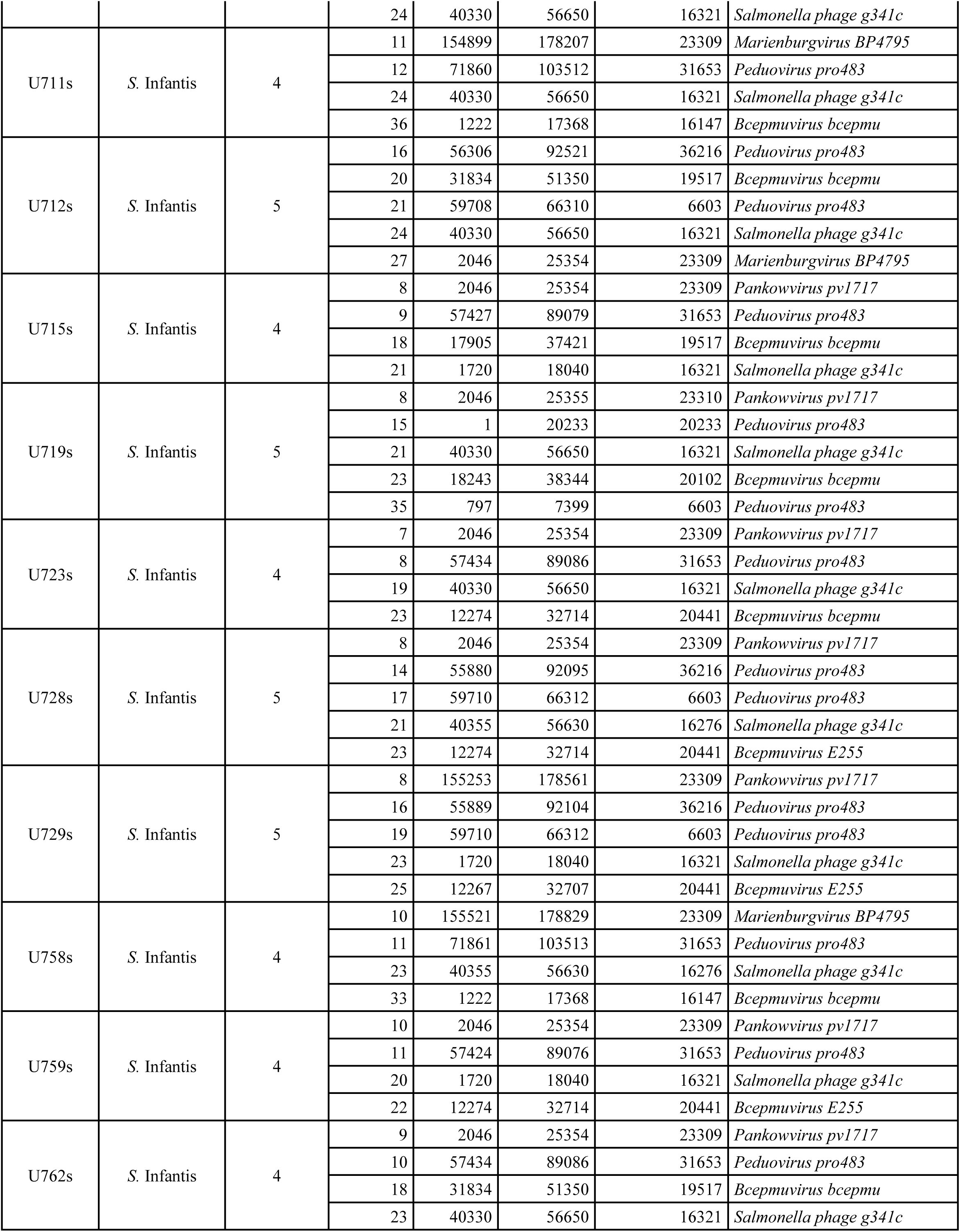
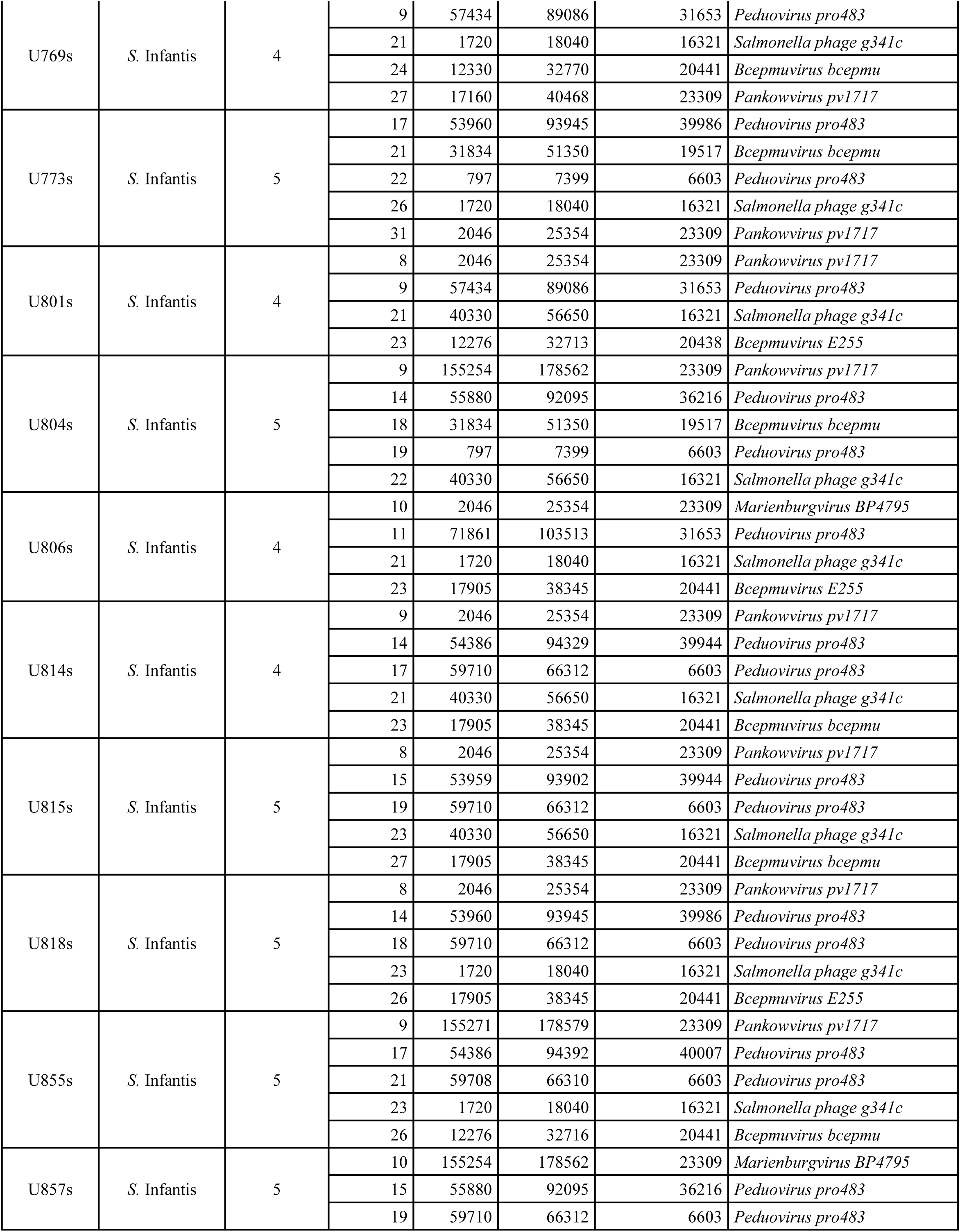
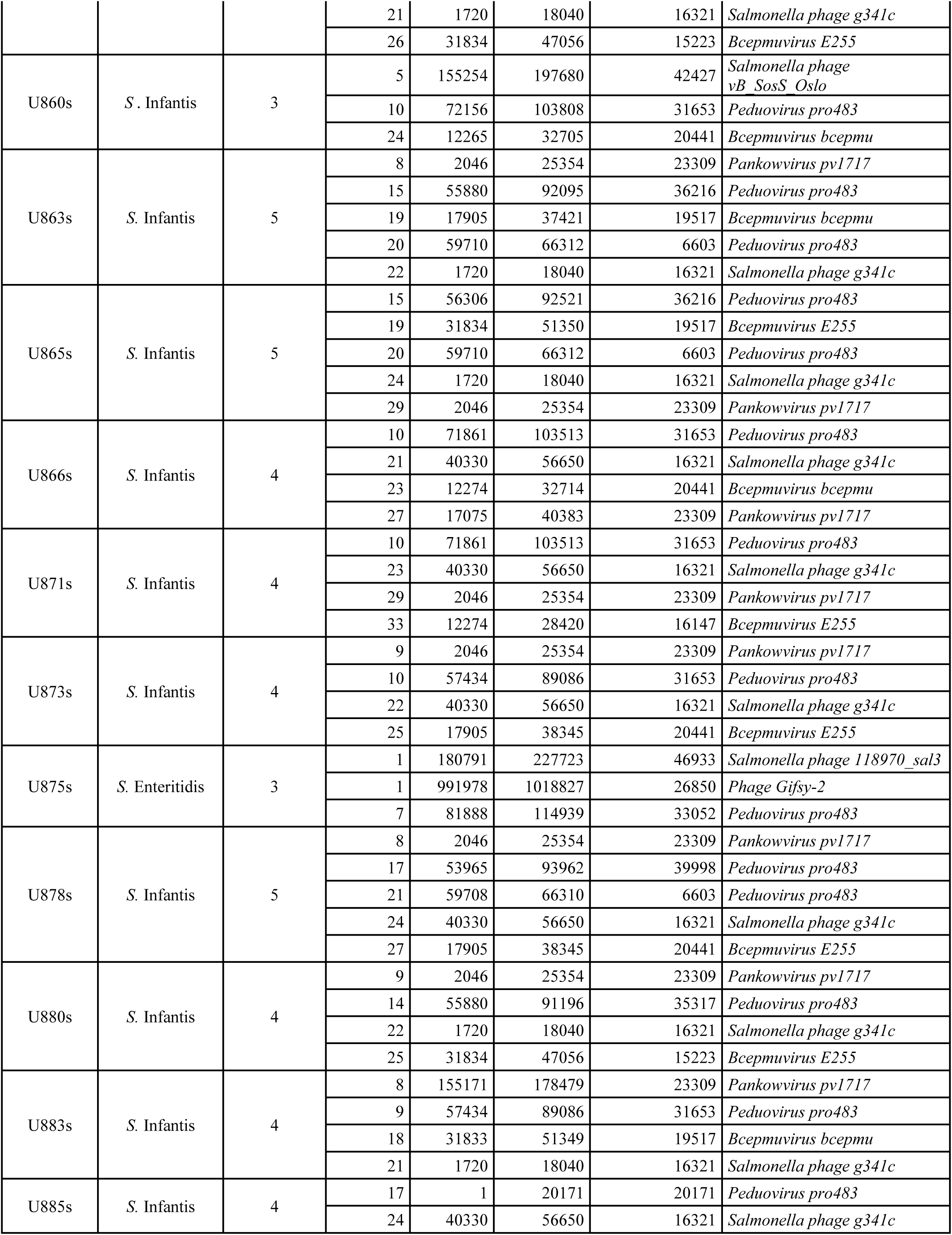
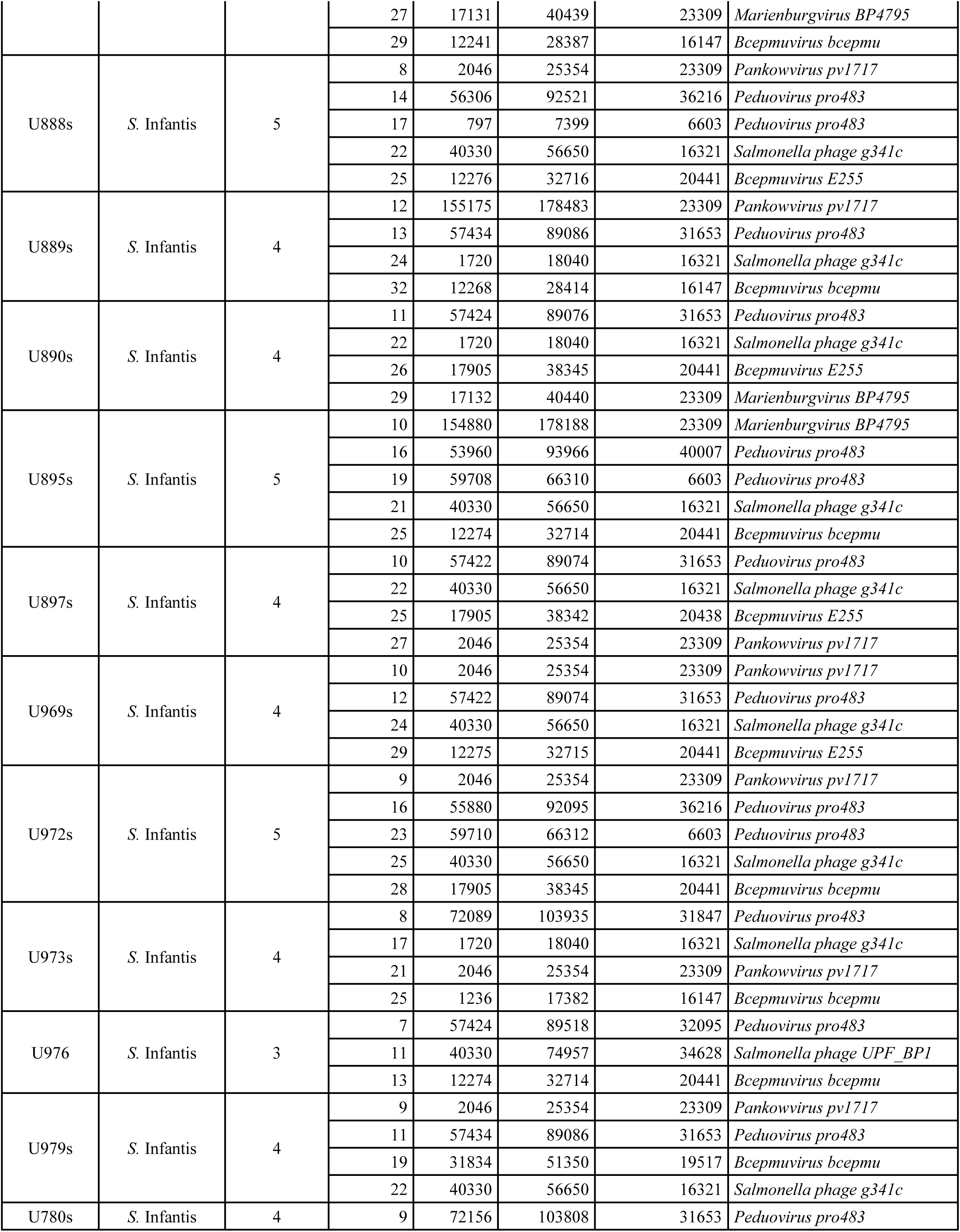
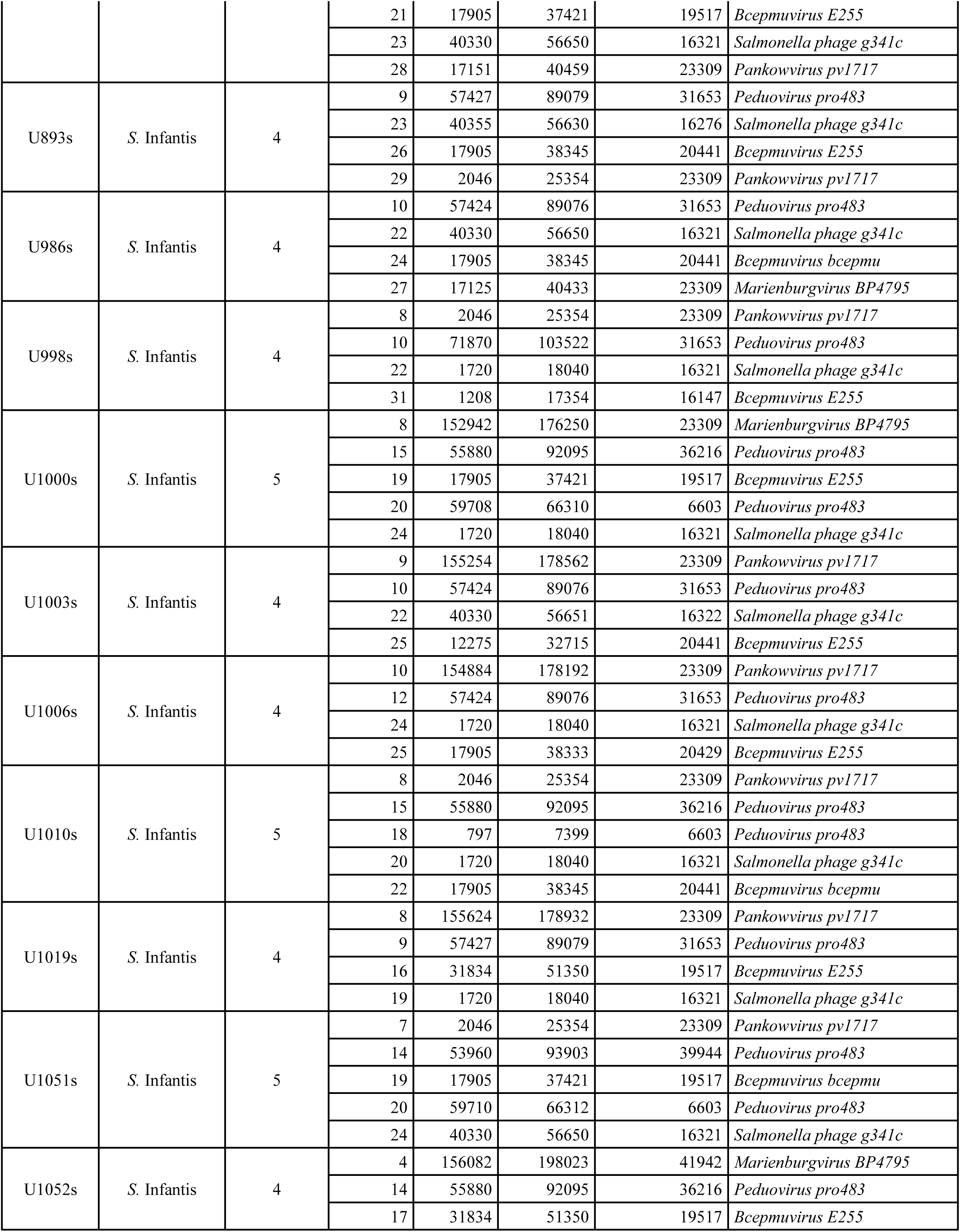
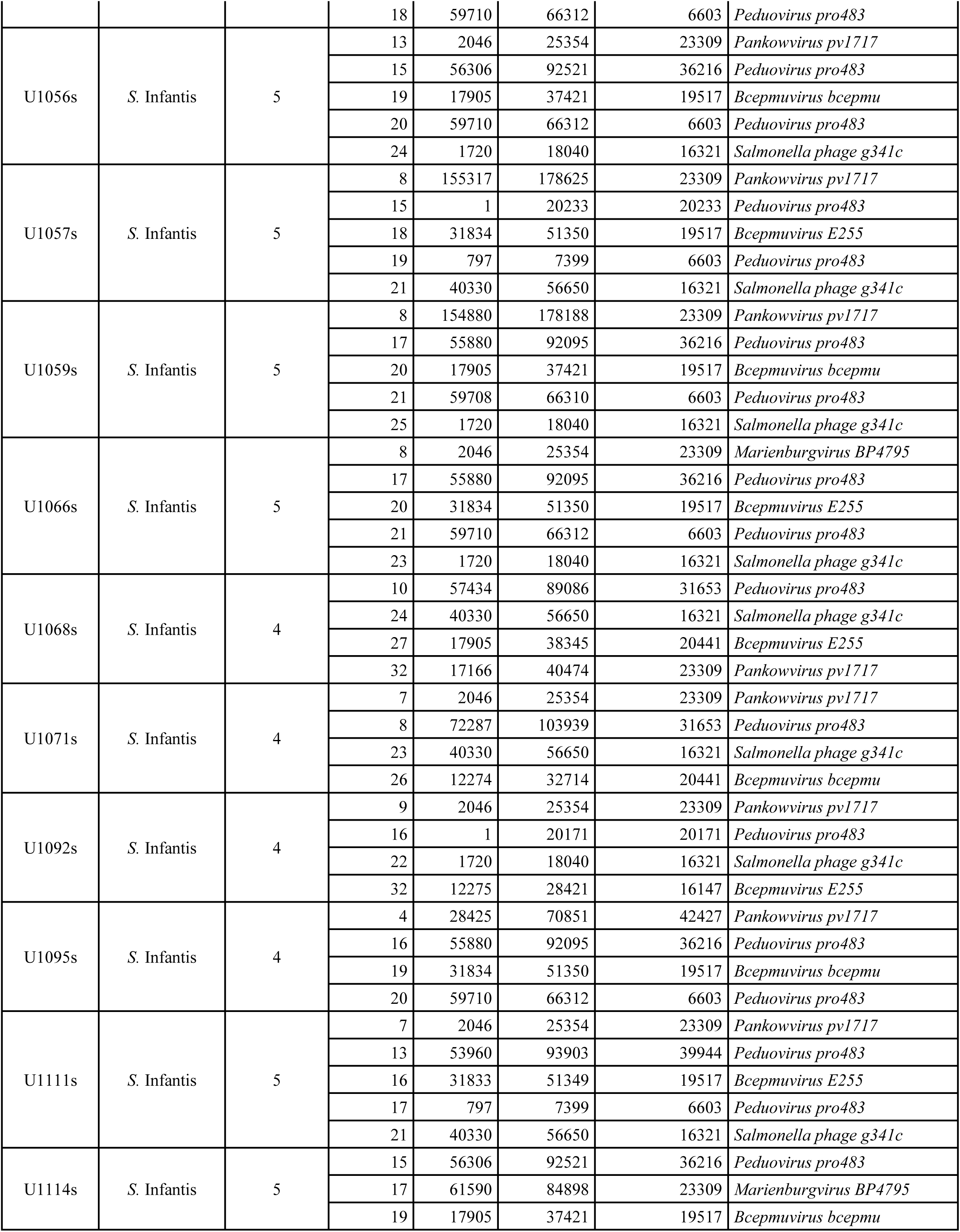
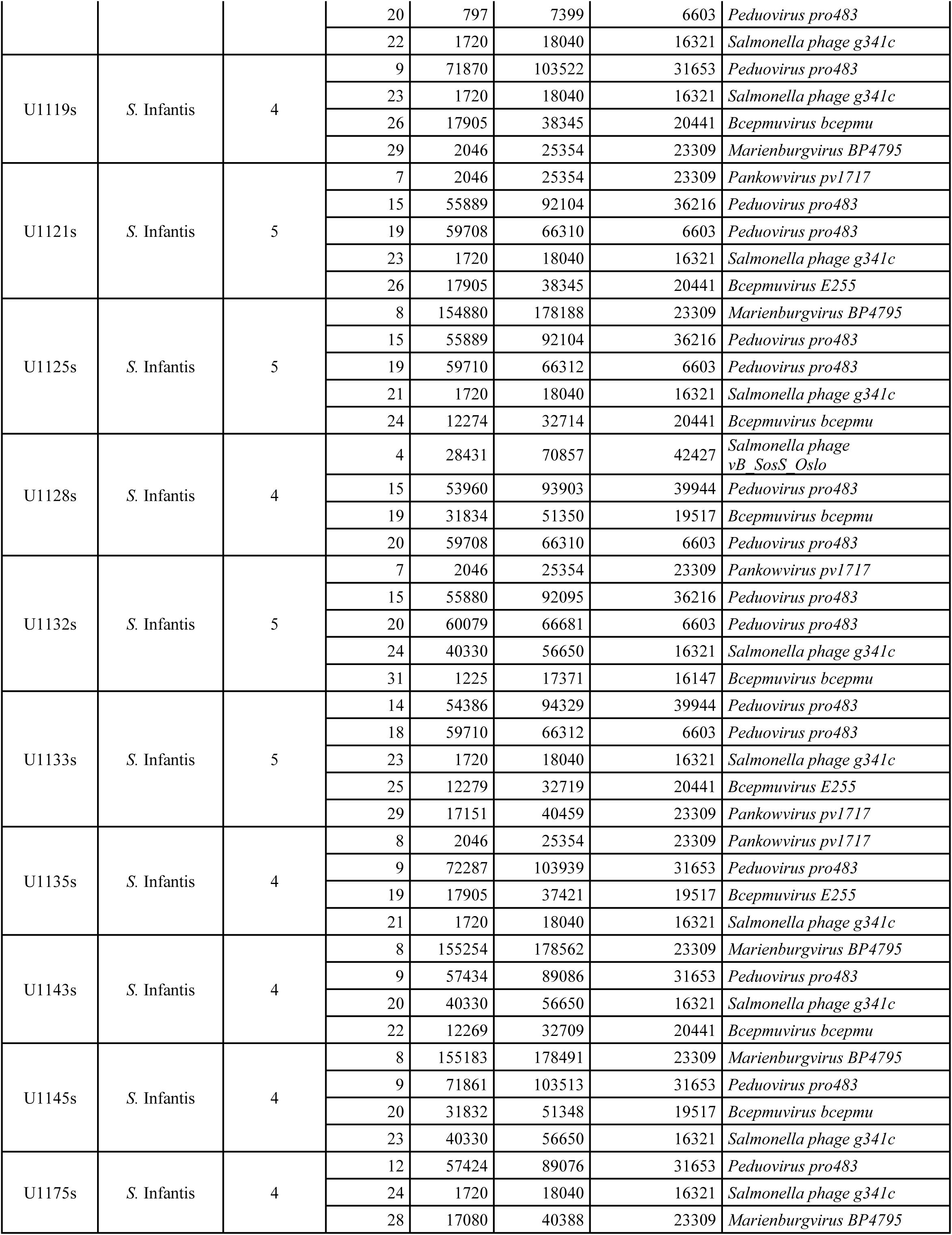
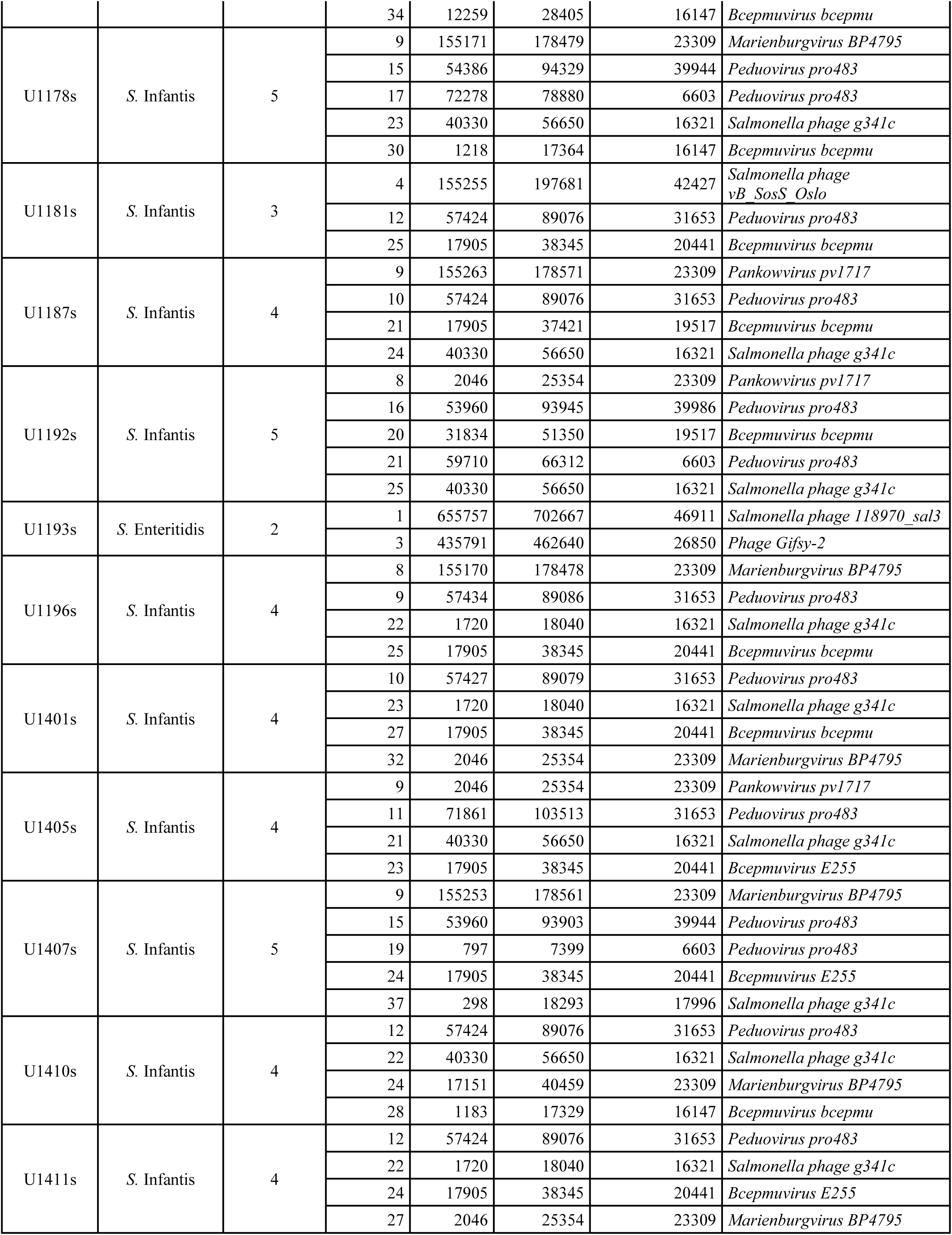
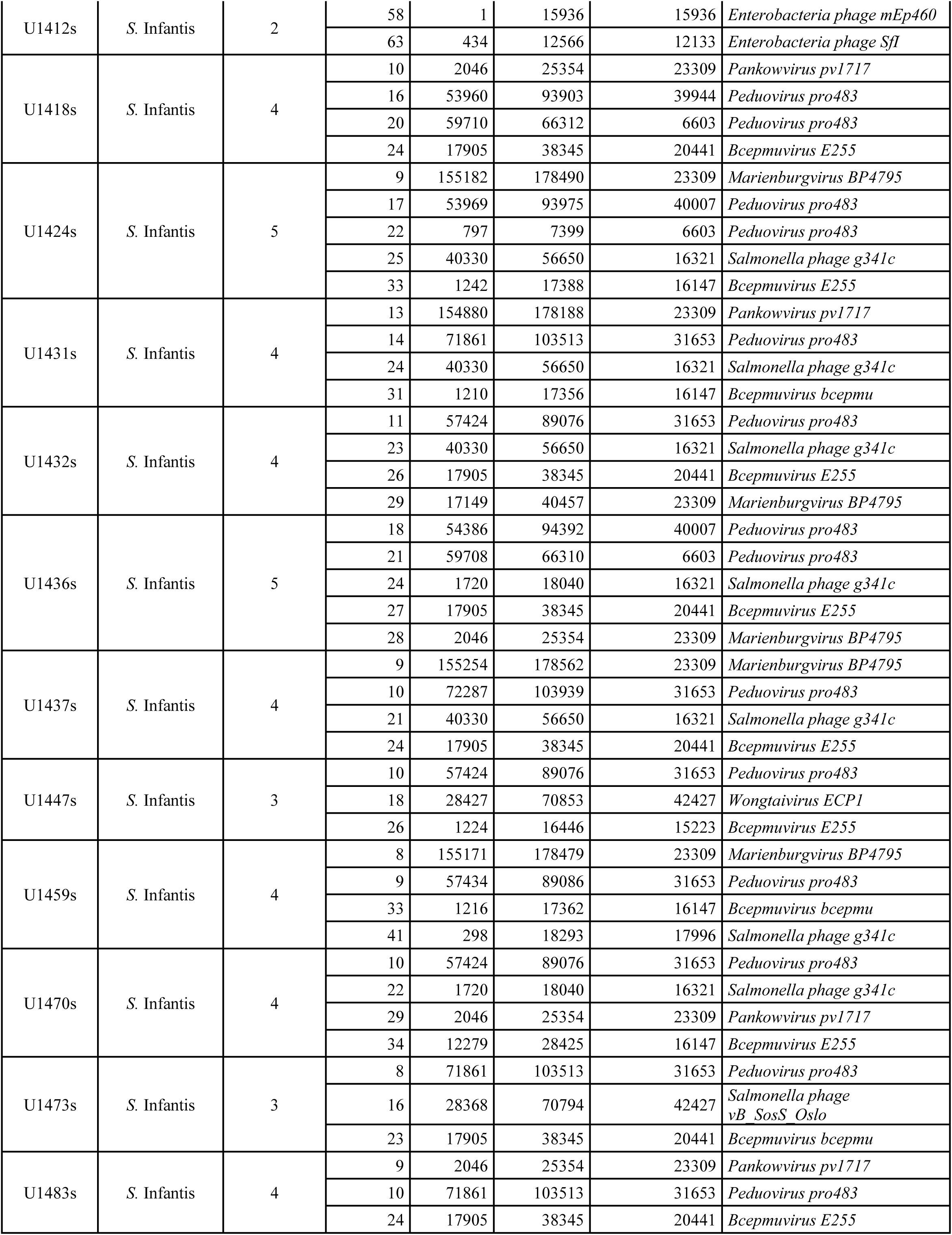
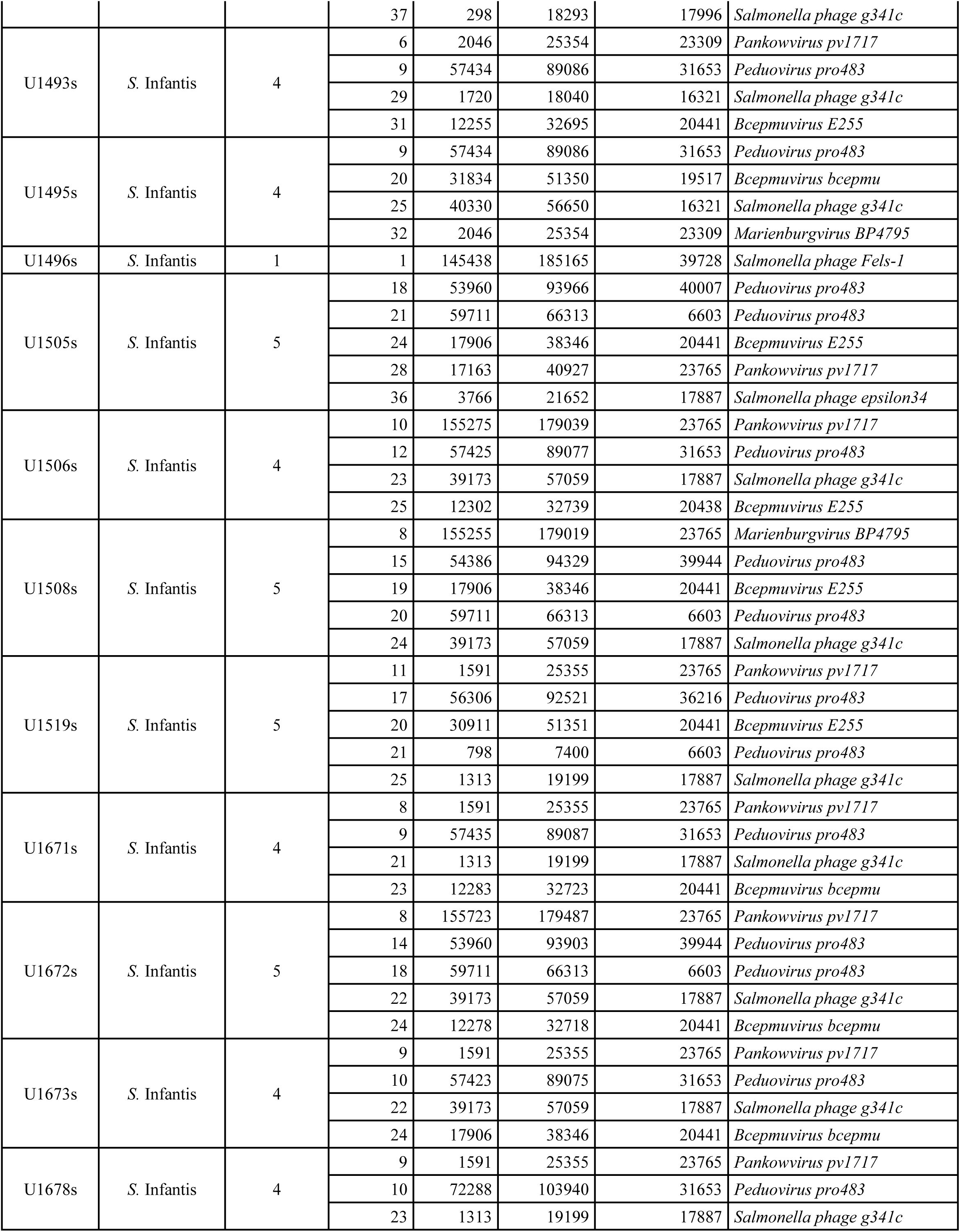
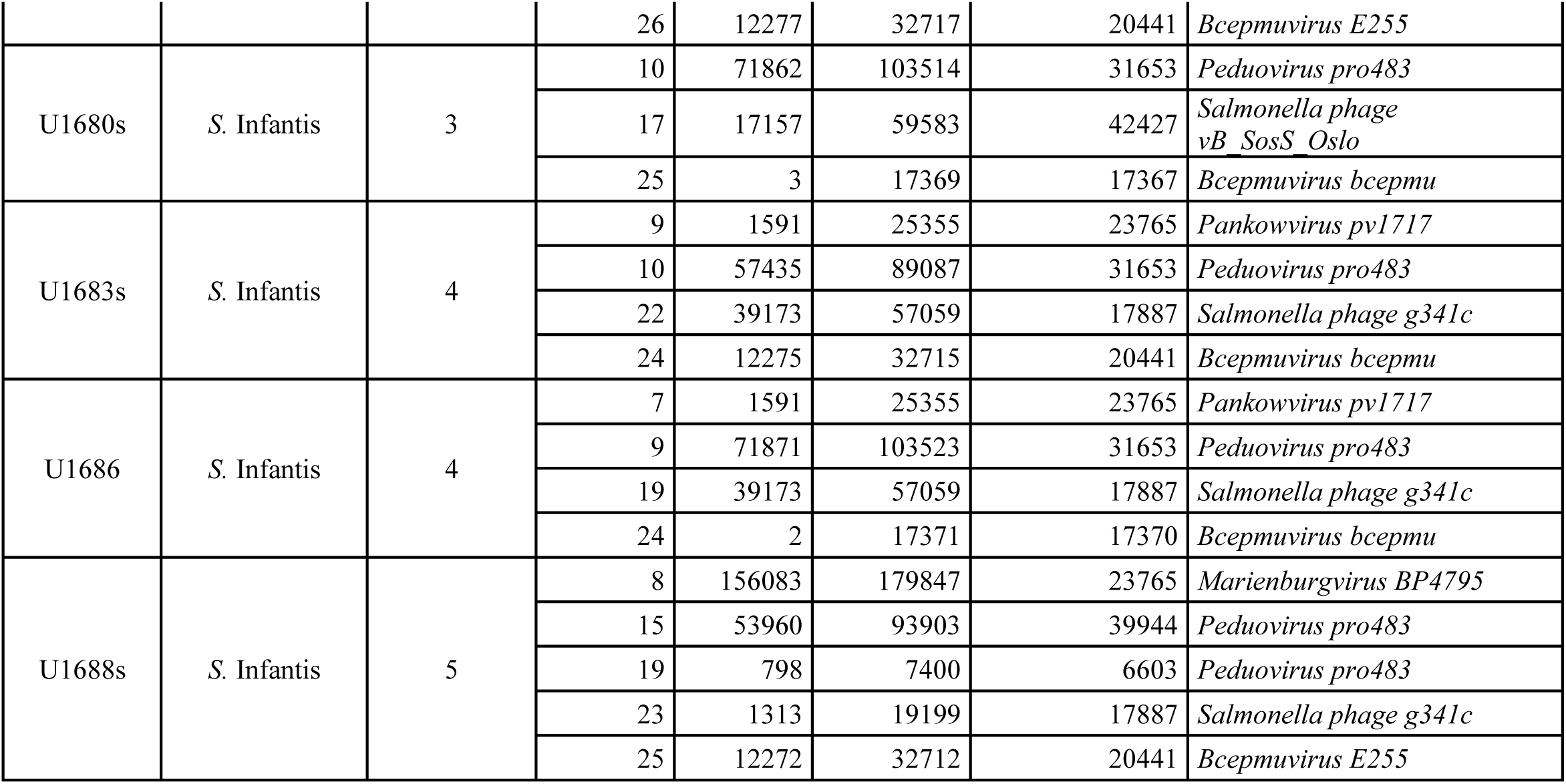
Prophages identified in the different S. enterica genomes analyzed, with their respective number of prophages per genome detected, scaffold, location and prophage length.

## References

1. Altschul, S. F., Gish, W., Miller, W., Myers, E. W., & Lipman, D. J. (1990). Basic local alignment search tool. Journal of Molecular Biology, 215(3), 403–410. 10.1016/S0022-2836(05)80360-2

2. Ammendola, S., Ajello, M., Pasquali, P., Kroll, J. S., Langford, P. R., Rotilio, G., Valenti, P., & Battistoni, A. (2005). Differential contribution of sodC1 and sodC2 to intracellular survival and pathogenicity of Salmonella enterica serovar Choleraesuis. Microbes and Infection, 7(4), 698–707. 10.1016/j.micinf.2005.01.005

3. Andrews, K., Landeryou, T., Sicheritz-Pontén, T., & Nale, J. Y. (2024). Diverse prophage elements of Salmonella enterica serovars show potential roles in bacterial pathogenicity. Cells, 13(6), 514.

4. Araya, D. V., Quiroz, T. S., Tobar, H. E., Lizana, R. J., Quezada, C. P., Santiviago, C. A., Riedel, C. A., Kalergis, A. M., & Bueno, S. M. (2010). Deletion of a prophage-like element causes attenuation of Salmonella enterica serovar Enteritidis and promotes protective immunity. Vaccine, 28(33), 5458–5466. 10.1016/j.vaccine.2010.05.073

5. Arndt, D., Marcu, A., Liang, Y., & Wishart, D. S. (2019). PHAST, PHASTER and PHASTEST: tools for finding prophage in bacterial genomes. Briefings in Bioinformatics, 20(4), 1560–1567.

6. Berngruber, T. W., Weissing, F. J., & Gandon, S. (2010). Inhibition of Superinfection and the Evolution of Viral Latency. Journal of Virology, 84(19), 10200–10208. 10.1128/jvi.00865-10

7. Bobay, L.-M., Touchon, M., & Rocha, E. P. C. (2014). Pervasive domestication of defective prophages by bacteria. Proceedings of the National Academy of Sciences, 111(33), 12127– 12132. 10.1073/pnas.1405336111

8. Bolger, A. M., Lohse, M., & Usadel, B. (2014). Trimmomatic: a flexible trimmer for Illumina sequence data. Bioinformatics, 30(15), 2114–2120.

9. Chen, Y., Liu, L., Guo, Y., Chu, J., Wang, B., Sui, Y., Wei, H., Hao, H., Huang, L., & Cheng, G. (2024). Distribution and genetic characterization of fluoroquinolone resistance gene qnr among Salmonella strains from chicken in China. Microbiology Spectrum, 12(4). 10.1128/spectrum.03000-23

10. Cohen, E., Rahav, G., & Gal-Mor, O. (2020). Genome Sequence of an Emerging Salmonella enterica Serovar Infantis and Genomic Comparison with Other S. Infantis Strains. Genome Biology and Evolution, 12(3), 223–228. 10.1093/gbe/evaa048

11. Colavecchio, A., D’Souza, Y., Tompkins, E., Jeukens, J., Freschi, L., Emond-Rheault, J., Kukavica-Ibrulj, I., Boyle, B., Bekal, S., Tamber, S., Levesque, R. C., & Goodridge, L. D. (2017). Prophage Integrase Typing Is a Useful Indicator of Genomic Diversity in Salmonella enterica. Frontiers in Microbiology, 8. 10.3389/fmicb.2017.01283

12. Coudert, E., Gehant, S., de Castro, E., Pozzato, M., Baratin, D., Neto, T., Sigrist, C. J. A., Redaschi, N., Bridge, A., & The UniProt Consortium. (2023). Annotation of biologically relevant ligands in UniProtKB using ChEBI. Bioinformatics, 39(1), btac793. 10.1093/bioinformatics/btac793

13. Crossland, W. L., Shaw, J. P., O’Leary, C., Gill, J., & Liu, M. (2019). Complete Genome Sequence of Citrobacter freundii Siphophage Sazh. Microbiology Resource Announcements, 8(50). 10.1128/mra.01317-19

14. D’Alessandro, B., Perez Escanda, V., Balestrazzi, L., Iriarte, A., Pickard, D., Yim, L., Chabalgoity, J. A., & Betancor, L. (2018). A novel prophage identified in strains from Salmonella enterica serovar Enteritidis is a phylogenetic signature of the lineage ST-1974. Microbial Genomics, 4(3), e000161.

15. Dion, M. B., Oechslin, F., & Moineau, S. (2020). Phage diversity, genomics and phylogeny. Nature Reviews Microbiology, 18(3), Article 3. 10.1038/s41579-019-0311-5

16. Dziva, F., Hauser, H., Connor, T. R., Van Diemen, P. M., Prescott, G., Langridge, G. C., Eckert, S., Chaudhuri, R. R., Ewers, C., Mellata, M., Mukhopadhyay, S., Curtiss, R., Dougan, G., Wieler, L. H., Thomson, N. R., Pickard, D. J., & Stevens, M. P. (2012). Sequencing and Functional Annotation of Avian Pathogenic Escherichia coli Serogroup O78 Strains Reveal the Evolution of E. coli Lineages Pathogenic for Poultry via Distinct Mechanisms. Infection and Immunity, 81(3), 838–849. 10.1128/iai.00585-12

17. Emmanuel, I. E. (2021). Challenges of Phage Therapy as a Strategic Tool for the Control of Salmonella Kentucky and Repertoire of Antibiotic Resistance Genes in Africa. In Bacteriophages in Therapeutics. IntechOpen. 10.5772/intechopen.95329

18. Ewels, P., Magnusson, M., Lundin, S., & Käller, M. (2016). MultiQC: summarize analysis results for multiple tools and samples in a single report. Bioinformatics, 32(19), 3047–3048. 10.1093/bioinformatics/btw354

19. Federici, S., Nobs, S. P., & Elinav, E. (2021). Phages and their potential to modulate the microbiome and immunity. Cellular & molecular immunology, 18(4), 889–904. 10.1038/s41423-020-00532-4

20. Figueroa-Bossi, N., & Bossi, L. (1999). Inducible prophages contribute to Salmonella virulence in mice. Molecular Microbiology, 33(1), 167–176. 10.1046/j.1365-2958.1999.01461.x

21. Foley, S. L., Johnson, T. J., Ricke, S. C., Nayak, R., & Danzeisen, J. (2013). Salmonella pathogenicity and host adaptation in Chicken-Associated serovars. Microbiology and Molecular Biology Reviews, 77(4), 582–607. 10.1128/mmbr.00015-13

22. Gao, R., Naushad, S., Moineau, S., Levesque, R., Goodridge, L., & Ogunremi, D. (2020). Comparative genomic analysis of 142 bacteriophages infecting Salmonella enterica subsp. Enterica. BMC Genomics, 21(1), 374. 10.1186/s12864-020-6765-z

23. Golubeva, Y. A., & Slauch, J. M. (2006). Salmonella enterica Serovar Typhimurium Periplasmic Superoxide Dismutase SodCI Is a Member of the PhoPQ Regulon and Is Induced in Macrophages. Journal of Bacteriology, 188(22), 7853–7861. 10.1128/JB.00706-06

24. Gorbalenya, A. E., & Lauber, C. (2022). Bioinformatics of virus taxonomy: foundations and tools for developing sequence-based hierarchical classification. Current Opinion in Virology, 52, 48–56

25. Gymoese, P., Kiil, K., Torpdahl, M., Østerlund, M. T., Sørensen, G., Olsen, J. E., Nielsen, E. M., & Litrup, E. (2019). WGS based study of the population structure of Salmonella enterica serovar Infantis. BMC Genomics, 20(1), 870. 10.1186/s12864-019-6260-6

26. Hatcher, E. L., Zhdanov, S. A., Bao, Y., Blinkova, O., Nawrocki, E. P., Ostapchuck, Y., … & Brister, J. R. (2017). Virus Variation Resource–improved response to emergent viral outbreaks. Nucleic acids research, 45(D1), D482–D490.

27. Hatfull, G. F., Jacobs-Sera, D., Lawrence, J. G., Pope, W. H., Russell, D. A., Ko, C.-C., Weber, R. J., Patel, M. C., Germane, K. L., Edgar, R. H., Hoyte, N. N., Bowman, C. A., Tantoco, A. T., Paladin, E. C., Myers, M. S., Smith, A. L., Grace, M. S., Pham, T. T., O’Brien, M. B., … Hendrix, R. W. (2010). Comparative Genomic Analysis of 60 Mycobacteriophage Genomes: Genome Clustering, Gene Acquisition, and Gene Size. Journal of Molecular Biology, 397(1), 119–143. 10.1016/j.jmb.2010.01.011

28. Henrot, C., & Petit, M.-A. (2022). Signals triggering prophage induction in the gut microbiota. Molecular Microbiology, 118(5), 494–502. 10.1111/mmi.14983

29. Ho, T. D., & Slauch, J. M. (2001). OmpC is the receptor for Gifsy-1 and Gifsy-2 bacteriophages of Salmonella. Journal of bacteriology, 183(4), 1495–1498. 10.1128/JB.183.4.1495-1498.2001

30. Holmes, E. C. (2011). What does virus evolution tell us about virus origins?. Journal of virology, 85(11), 5247–5251

31. Huehn, S., La Ragione, R. M., Anjum, M., Saunders, M., Woodward, M. J., Bunge, C., Helmuth, R., Hauser, E., Guerra, B., Beutlich, J., Brisabois, A., Peters, T., Svensson, L., Madajczak, G., Litrup, E., Imre, A., Herrera-Leon, S., Mevius, D., Newell, D. G., & Malorny, B. (2009). Virulotyping and antimicrobial resistance typing ofSalmonella entericaSerovars relevant to human health in Europe. Foodborne Pathogens and Disease, 7(5), 523–535. 10.1089/fpd.2009.0447

32. Hyatt, D., Chen, G.-L., LoCascio, P. F., Land, M. L., Larimer, F. W., & Hauser, L. J. (2010). Prodigal: Prokaryotic gene recognition and translation initiation site identification. BMC Bioinformatics, 11(1), 119. 10.1186/1471-2105-11-119

33. Jackman, S. D., Vandervalk, B. P., Mohamadi, H., Chu, J., Yeo, S., Hammond, S. A., Jahesh, G., Khan, H., Coombe, L., Warren, R. L., & Birol, I. (2017). ABySS 2.0: Resource-efficient assembly of large genomes using a Bloom filter. Genome Research, 27(5), 768– 777. 10.1101/gr.214346.116

34. Jonge, P. A. de, Nobrega, F. L., Brouns, S. J. J., & Dutilh, B. E. (2019). Molecular and Evolutionary Determinants of Bacteriophage Host Range. Trends in Microbiology, 27(1), 51–63. 10.1016/j.tim.2018.08.006

35. Katoh, K., Rozewicki, J., & Yamada, K. D. (2019). MAFFT online service: Multiple sequence alignment, interactive sequence choice and visualization. Briefings in Bioinformatics, 20(4), 1160–1166. 10.1093/bib/bbx108

36. Kearse, M., Moir, R., Wilson, A., Stones-Havas, S., Cheung, M., Sturrock, S., Buxton, S., Cooper, A., Markowitz, S., Duran, C., Thierer, T., Ashton, B., Meintjes, P., & Drummond, A. (2012). Geneious Basic: An integrated and extendable desktop software platform for the organization and analysis of sequence data. Bioinformatics (Oxford, England), 28(12), 1647–1649. 10.1093/bioinformatics/bts199

37. Kirkwood, M., Vohra, P., Bawn, M., Thilliez, G., Pye, H., Tanner, J., Chintoan-Uta, C., Branchu, P., Petrovska, L., Dallman, T., Hall, N., Stevens, M. P., & Kingsley, R. A. (2021). Ecological niche adaptation of Salmonella Typhimurium U288 is associated with altered pathogenicity and reduced zoonotic potential. Communications Biology, 4(1), Article 1. 10.1038/s42003-021-02013-4

38. Kondo, K., Kawano, M., & Sugai, M. (2021). Distribution of Antimicrobial Resistance and Virulence Genes within the Prophage-Associated Regions in Nosocomial Pathogens. mSphere, 6(4). 10.1128/msphere.00452-21

39. Kongari, R., Rajaure, M., Cahill, J., Rasche, E., Mijalis, E., Berry, J., & Young, R. (2018). Phage spanins: diversity, topological dynamics and gene convergence. BMC Bioinformatics, 19(1). 10.1186/s12859-018-2342-8

40. Koonjan, S., Seijsing, F., Cooper, C. J., & Nilsson, A. S. (2020). Infection Kinetics and phylogenetic analysis of VB_ECOD_SU57, a virulent T1-Like drexlerviridae coliphage. Frontiers in Microbiology, 11. 10.3389/fmicb.2020.565556

41. Ksibi, B., Ktari, S., Ghedira, K., Othman, H., Maalej, S., Mnif, B., Fabre, L., Rhimi, F., Hello, S. L., & Hammami, A. (2022). Antimicrobial resistance genes, virulence markers and prophage sequences in Salmonella enterica serovar Enteritidis isolated in Tunisia using whole genome sequencing. Current Research in Microbial Sciences, 3, 100151. 10.1016/j.crmicr.2022.100151

42. Kürekci, C., Sahin, S., Iwan, E., Kwit, R., Bomba, A., & Wasyl, D. (2020). Whole-genome sequence analysis of Salmonella Infantis isolated from raw chicken meat samples and insights into pESI-like megaplasmid. International Journal of Food Microbiology, 337, 108956. 10.1016/j.ijfoodmicro.2020.108956

43. Lee, W. W. Y., Mattock, J., Greig, D. R., Langridge, G. C., Baker, D., Bloomfield, S., Mather, A. E., Wain, J. R., Edwards, A. M., Hartman, H., Dallman, T. J., Chattaway, M. A., & Nair, S. (2021). Characterization of a pESI-like plasmid and analysis of multidrug-resistant Salmonella enterica Infantis isolates in England and Wales. Microbial Genomics, 7(10). 10.1099/mgen.0.000658

44. Lima-Mendez, G., Van Helden, J., Toussaint, A., & Leplae, R. (2008). Reticulate Representation of Evolutionary and Functional Relationships between Phage Genomes. Molecular Biology and Evolution, 25(4), 762–777. 10.1093/molbev/msn023

45. Liu, Z., Wang, L., Yu, Y., Fotin, A., Wang, Q., Gao, P., Zhang, Y., Fotina, T., & Ma, J. (2022). STEE enhances the virulence of salmonella pullorum in chickens by regulating the inflammation response. Frontiers in Veterinary Science, 9. 10.3389/fvets.2022.926505

46. Maffei, E., Shaidullina, A., Burkolter, M., Heyer, Y., Estermann, F., Druelle, V., Sauer, P., Willi, L., Michaelis, S., Hilbi, H., Thaler, D. S., & Harms, A. (2021). Systematic exploration of Escherichia coli phage-host interactions with the BASEL phage collection. PLoS biology, 19(11), e3001424. 10.1371/journal.pbio.3001424

47. Malberg Tetzschner, A. M., Johnson, J. R., Johnston, B. D., Lund, O., & Scheutz, F. (2020). In silico genotyping of Escherichia coli isolates for extraintestinal virulence genes by use of whole-genome sequencing data. Journal of clinical microbiology, 58(10), 10–1128.

48. Manni, M., Berkeley, M. R., Seppey, M., Simão, F. A., & Zdobnov, E. M. (2021). BUSCO Update: Novel and Streamlined Workflows along with Broader and Deeper Phylogenetic Coverage for Scoring of Eukaryotic, Prokaryotic, and Viral Genomes. Molecular Biology and Evolution, 38(10), 4647–4654. 10.1093/molbev/msab199

49. Mejía, L., Medina, J. L., Bayas, R., Salazar, C. S., Villavicencio, F., Zapata, S., Matheu, J., Wagenaar, J. A., González-Candelas, F., & Vinueza-Burgos, C. (2020). Genomic Epidemiology of Salmonella Infantis in Ecuador: From Poultry Farms to Human Infections. Frontiers in veterinary science, 7, 547891. 10.3389/fvets.2020.547891

50. Mohammed, M., Casjens, S. R., Millard, A. D., Harrison, C., Gannon, L., & Chattaway, M. A. (2023). Genomic analysis of Anderson typing phages of Salmonella Typhimrium: Towards understanding the basis of bacteria-phage interaction. Scientific Reports, 13(1), Article 1. 10.1038/s41598-023-37307-6

51. Monteiro, R., Pires, D. P., Costa, A. R., & Azeredo, J. (2019). Phage Therapy: Going Temperate? Trends in Microbiology, 27(4), 368–378. 10.1016/j.tim.2018.10.008

52. Montoro-Dasi, L., Lorenzo-Rebenaque, L., Marco-Fuertes, A., Vega, S., & Marin, C. (2023). Holistic Strategies to Control Salmonella Infantis: An Emerging Challenge in the European Broiler Sector. Microorganisms, 11(7), Article 7. 10.3390/microorganisms11071765

53. Morgulis, A., Coulouris, G., Raytselis, Y., Madden, T. L., Agarwala, R., & Schäffer, A. A. (2008). Database indexing for production MegaBLAST searches. Bioinformatics (Oxford, England), 24(16), 1757–1764. 10.1093/bioinformatics/btn322

54. Mottawea, W., Duceppe, M.-O., Dupras, A. A., Usongo, V., Jeukens, J., Freschi, L., Emond-Rheault, J.-G., Hamel, J., Kukavica-Ibrulj, I., Boyle, B., Gill, A., Burnett, E., Franz, E., Arya, G., Weadge, J. T., Gruenheid, S., Wiedmann, M., Huang, H., Daigle, F., … Ogunremi, D. (2018). Salmonella enterica Prophage Sequence Profiles Reflect Genome Diversity and Can Be Used for High Discrimination Subtyping. Frontiers in Microbiology, 9, 836. 10.3389/fmicb.2018.00836

55. Moura de Sousa, J. A., Pfeifer, E., Touchon, M., & Rocha, E. P. C. (2021). Causes and Consequences of Bacteriophage Diversification via Genetic Exchanges across Lifestyles and Bacterial Taxa. Molecular Biology and Evolution, 38(6), 2497–2512. 10.1093/molbev/msab044

56. Nayfach, S., Camargo, A. P., Schulz, F., Eloe-Fadrosh, E., Roux, S., & Kyrpides, N. C. (2021). CheckV assesses the quality and completeness of metagenome-assembled viral genomes. Nature biotechnology, 39(5), 578–585.

57. Nishijima, S., Nagata, N., Kiguchi, Y., Kojima, Y., Miyoshi-Akiyama, T., Kimura, M., Ohsugi, M., Ueki, K., Oka, S., Mizokami, M., Itoi, T., Kawai, T., Uemura, N., & Hattori, M. (2022). Extensive gut virome variation and its associations with host and environmental factors in a population-level cohort. Nature Communications, 13(1), Article 1. 10.1038/s41467-022-32832-w

58. Nishimura, Y., Yamada, K., Okazaki, Y., & Ogata, H. (2024). DiGAlign: Versatile and Interactive Visualization of Sequence Alignment for Comparative Genomics. Microbes and environments, 39(1), ME23061.

59. Owen, S. V., Canals, R., Wenner, N., Hammarlöf, D. L., Kröger, C., & Hinton, J. C. D. (2020). A window into lysogeny: revealing temperate phage biology with transcriptomics. Microbial Genomics, 6(2). 10.1099/mgen.0.000330

60. Owen, S. V., Wenner, N., Dulberger, C. L., Rodwell, E. V., Bowers-Barnard, A., Quinones-Olvera, N., Rigden, D. J., Rubin, E. J., Garner, E. C., Baym, M., & Hinton, J. C. D. (2021). Prophages encode phage-defense systems with cognate self-immunity. Cell Host & Microbe, 29(11), 1620–1633.e8. 10.1016/j.chom.2021.09.002

61. Pacífico, C., Hilbert, M., Sofka, D., Dinhopl, N., Pap, I., Aspöck, C., Carriço, J. A., & Hilbert, F. (2019). Natural occurrence of Escherichia coli-Infecting bacteriophages in clinical samples. Frontiers in Microbiology, 10. 10.3389/fmicb.2019.02484

62. Parker, C. T., Huynh, S., Alexander, A., Oliver, A. S., & Cooper, K. K. (2021). Genomic Characterization of Salmonella typhimurium DT104 Strains Associated with Cattle and Beef Products. Pathogens, 10(5), Article 5. 10.3390/pathogens10050529

63. Parra, M., De Los Ángeles Bayas-Rea, R., Guerrero, T., Torres, S., D’Auria, G., Reyes-Prieto, M., & Zapata, S. (2023). Complete Genome Sequences of Two Lytic Phages of Salmonella enterica Isolated from Wastewater in Ecuador. Microbiology Resource Announcements, 12(2). 10.1128/mra.01048-22

64. Petrovska, L., Mather, A. E., AbuOun, M., Branchu, P., Harris, S. R., Connor, T., Hopkins, K. L., Underwood, A., Lettini, A. A., & Page, A. (2016). Microevolution of monophasic Salmonella typhimurium during epidemic, United Kingdom, 2005–2010. Emerging Infectious Diseases, 22(4), 617.

65. Pope, W. H., Jacobs-Sera, D., Russell, D. A., Peebles, C. L., Al-Atrache, Z., Alcoser, T. A., Alexander, L. M., Alfano, M. B., Alford, S. T., Amy, N. E., Anderson, M. D., Anderson, A. G., Ang, A. A. S., Ares, M., Barber, A. J., Barker, L. P., Barrett, J. M., Barshop, W. D., Bauerle, C. M., … Hatfull, G. F. (2011). Expanding the Diversity of Mycobacteriophages: Insights into Genome Architecture and Evolution. PLoS ONE, 6(1), e16329. 10.1371/journal.pone.0016329

66. Potter, S. C., Luciani, A., Eddy, S. R., Park, Y., Lopez, R., & Finn, R. D. (2018). HMMER web server: 2018 update. Nucleic acids research, 46(W1), W200–W204.

67. Prjibelski, A., Antipov, D., Meleshko, D., Lapidus, A., & Korobeynikov, A. (2020). Using SPAdes De Novo Assembler. Current Protocols in Bioinformatics, 70(1), e102. 10.1002/cpbi.102

68. Rusconi, B., Sanjar, F., Koenig, S. S. K., Mammel, M. K., Tarr, P. I., & Eppinger, M. (2016). Whole Genome Sequencing for Genomics-Guided Investigations of Escherichia coli O157:H7 Outbreaks. Frontiers in Microbiology, 7. https://www.frontiersin.org/articles/10.3389/fmicb.2016.00985

69. Sayers, E. W., Bolton, E. E., Brister, J. R., Canese, K., Chan, J., Comeau, D. C., Connor, R., Funk, K., Kelly, C., Kim, S., Madej, T., Marchler-Bauer, A., Lanczycki, C., Lathrop, S., Lu, Z., Thibaud-Nissen, F., Murphy, T., Phan, L., Skripchenko, Y., … Sherry, S. T. (2022). Database resources of the national center for biotechnology information. Nucleic Acids Research, 50(D1), D20–D26. 10.1093/nar/gkab1112

70. Schwengers, O., Jelonek, L., Dieckmann, M. A., Beyvers, S., Blom, J., & Goesmann, A. (2021). Bakta: Rapid and standardized annotation of bacterial genomes via alignment-free sequence identification. Microbial Genomics, 7(11), 000685. 10.1099/mgen.0.000685

71. Shridhar, P. B., Worley, J. N., Gao, X., Yang, X., Noll, L. W., Shi, X., Bai, J., Meng, J., & Nagaraja, T. G. (2019). Analysis of virulence potential of Escherichia coli O145 isolated from cattle feces and hide samples based on whole genome sequencing. PLOS ONE, 14(11), e0225057. 10.1371/journal.pone.0225057

72. Starikova, E. V., Tikhonova, P. O., Prianichnikov, N. A., Rands, C. M., Zdobnov, E. M., Ilina, E. N., & Govorun, V. M. (2020). Phigaro: high-throughput prophage sequence annotation. Bioinformatics, 36(12), 3882–3884.

73. Switt, A. I. M., Sulakvelidze, A., Wiedmann, M., Kropinski, A. M., Wishart, D. S., Poppe, C., & Liang, Y. (2015). Salmonella Phages and Prophages: Genomics, Taxonomy, and Applied Aspects. In H. Schatten & A. Eisenstark (Eds.), Salmonella (Vol. 1225, pp. 237–287). Springer New York. 10.1007/978-1-4939-1625-2_15

74. Tisza, M. J., & Buck, C. B. (2021). A catalog of tens of thousands of viruses from human metagenomes reveals hidden associations with chronic diseases. Proceedings of the National Academy of Sciences, 118(23), e2023202118. 10.1073/pnas.2023202118

75. Turner, D., Kropinski, A. M., & Adriaenssens, E. M. (2021). A Roadmap for Genome-Based Phage Taxonomy. Viruses, 13(3), 506. 10.3390/v13030506

76. Vaid, R. K., Thakur, Z., Anand, T., Kumar, S., & Tripathi, B. N. (2021). Comparative genome analysis of Salmonella enterica serovar Gallinarum biovars Pullorum and Gallinarum decodes strain specific genes. PLoS ONE, 16(8), e0255612. 10.1371/journal.pone.0255612

77. Vallejos-Sánchez, K., Tataje-Lavanda, L., Villanueva-Pérez, D., Bendezú, J., Montalván, Á., Zimic-Peralta, M., Fernández-Sánchez, M., & Fernández-Díaz, M. (2019). Whole-Genome Sequencing of a Salmonella enterica subsp. Enterica Serovar Infantis Strain Isolated from Broiler Chicken in Peru. Microbiology Resource Announcements, 8(43), 10.1128/mra.00826-19. 10.1128/mra.00826-19

78. Vinueza-Burgos, C., Baquero, M., Medina, J., & De Zutter, L. (2019). Occurrence, genotypes and antimicrobial susceptibility of Salmonella collected from the broiler production chain within an integrated poultry company. International Journal of Food Microbiology, 299, 1–7. 10.1016/j.ijfoodmicro.2019.03.014

79. Wahl, A., Battesti, A., & Ansaldi, M. (2019). Prophages in Salmonella enterica: A driving force in reshaping the genome and physiology of their bacterial host? Molecular Microbiology, 111(2), 303–316. 10.1111/mmi.14167

80. Walker, P. J., Siddell, S. G., Lefkowitz, E. J., Mushegian, A. R., Adriaenssens, E. M., Dempsey, D. M., Dutilh, B. E., Harrach, B., Harrison, R. L., Hendrickson, R. C., Junglen, S., Knowles, N. J., Kropinski, A. M., Krupovic, M., Kuhn, J. H., Nibert, M., Orton, R. J., Rubino, L., Sabanadzovic, S., … Davison, A. J. (2020). Changes to virus taxonomy and the Statutes ratified by the International Committee on Taxonomy of Viruses (2020). Archives of Virology, 165(11), 2737–2748. 10.1007/s00705-020-04752-x

81. Weissman, J. L., Holmes, R., Barrangou, R., Moineau, S., Fagan, W. F., Levin, B., & Johnson, P. L. F. (2018). Immune loss as a driver of coexistence during host-phage coevolution. The ISME Journal, 12(2), Article 2. 10.1038/ismej.2017.194

82. Wendling, C. C. (2023). Prophage mediated control of higher order interactions—Insights from multi-level approaches. Current Opinion in Systems Biology, 35, 100469. 10.1016/j.coisb.2023.100469

83. Yu, P., Mathieu, J., Yang, Y., & Alvarez, P. J. J. (2017). Suppression of Enteric Bacteria by Bacteriophages: Importance of Phage Polyvalence in the Presence of Soil Bacteria. Environmental Science & Technology, 51(9), 5270–5278. 10.1021/acs.est.7b00529

